# Proteasome-dependent truncation of the negative heterochromatin regulator Epe1 mediates antifungal resistance

**DOI:** 10.1101/2021.12.20.473483

**Authors:** Imtiyaz Yaseen, Sharon A. White, Sito Torres-Garcia, Christos Spanos, Marcel Lafos, Elisabeth Gaberdiel, Rebecca Yeboah, Meriem El Karoui, Juri Rappsilber, Alison L. Pidoux, Robin C. Allshire

## Abstract

Epe1 histone demethylase restricts H3K9-methylation-dependent heterochromatin, preventing it from spreading over, and silencing, gene-containing regions in fission yeast. External stress induces an adaptive response allowing heterochromatin island formation that confers resistance on surviving wild-type lineages. Here we investigate the mechanism by which Epe1 is regulated in response to stress. Exposure to caffeine or antifungals results in Epe1 ubiquitylation and proteasome-dependent removal of the N-terminal 150 residues from Epe1, generating truncated tEpe1 which accumulates in the cytoplasm. Constitutive tEpe1 expression increases H3K9 methylation over several chromosomal regions, reducing expression of underlying genes and enhancing resistance. Reciprocally, constitutive non-cleavable Epe1 expression decreases resistance. tEpe1-mediated resistance requires a functional JmjC demethylase domain. Moreover, caffeine-induced Epe1-to-tEpe1 cleavage is dependent on an intact cell-integrity MAP kinase stress signalling pathway, mutations in which alter resistance. Thus, environmental changes provoke a mechanism that curtails the function of this key epigenetic modifier, allowing heterochromatin to reprogram gene expression, thereby bestowing resistance to some cells within a population. H3K9me-heterochromatin components are conserved in human and crop plant fungal pathogens for which a limited number of antifungals exist. Our findings reveal how transient heterochromatin-dependent antifungal resistant epimutations develop and thus inform on how they might be countered.

## Main

The overuse in agriculture of antifungal agents, related to those used in treating human fungal infections, has caused progressive increases in resistance in soil-borne fungi^1^. Consequently, clinical treatment of patients exhibiting aspergillosis (*Aspergillus*), candidiasis (*Candida*) or cryptococcosis (*Cryptococcus*) is challenging due to the limited number of effective antifungal drugs and the increasing prevalence of resistance^2–4^. A dilemma exists as widespread use of antifungals in agriculture is required to combat major plant pathogens such as *Magnaporthe oryzae* (rice; blast fungus) and *Zymoseptoria tritici* (wheat; leaf blotch) in order to enhance crop yields. Optimised crop production is required to meet the nutritional needs of the human population that is estimated to increase by approximately 2 billion in the next 30 years^5^. It is therefore important to understand the mechanisms that allow fungi to adapt and develop resistance to external insults such as antifungals in order to design more prudent interventions in clinical and agricultural settings.

Throughout their evolutionary history fungi have been exposed to challenging environments^6^. Consequently, fungi have developed robust, intricate systems to sense and adapt to ‘new’ external insults such as antifungal compounds. Altered environments or manipulations are known to allow transcriptional memory and epigenetic inheritance in single-celled organisms such as fungi^7–13^, and also in multicellular organisms^14–17^ so that resulting DNA and chromatin modifications can persist through cell division, long after the original stimulus has dissipated. Such ‘epigenetic memory’ encourages heterogeneity with variable persistence in otherwise genetically identical cell populations, providing a bet-hedging strategy that ensures adaptation and survival of a proportion of individual cells upon exposure to new environmental challenges. Thus, transient but metastable epigenetic states can confer a selective advantage to particular cell lineages within a clonal population, while allowing return to the initial normal state once external pressures are relaxed^13^.

The fission yeast Clr4 methyltransferase installs all histone H3K9 methylation, triggering heterochromatin formation and transcriptional repression at locations such as centromeres, telomeres, the mating-type locus^18–21^, and facultative heterochromatin islands^22–26^. Heterochromatin assembly is antagonized by the histone acetyltransferase Mst2, which acetylates H3, preventing H3K9 methylation, and the JmjC domain histone demethylase Epe1 that likely removes superfluous H3K9 methylation from euchromatic regions and islands^22, 27–30^. *S. pombe* lacks DNA methylation^31, 32^, thus its heterochromatin-mediated epigenetic regulation relies entirely on histone modification.

In the absence of Epe1 synthetic heterochromatin induced at ectopic chromosomal locations can be transmitted through multiple cell divisions by a read-write mechanism following the release of the heterochromatin-nucleating activity^7, 8^. Recently we demonstrated that heterochromatin redistributes in an acute response to caffeine stress^13^. Caffeine treatment resulted in Epe1 down-regulation, suggesting that this allows heterochromatin assembly at beneficial loci. However, the mechanism of Epe1 regulation in response to caffeine stress remained unexplained.

Here we determine how Epe1 levels are down-regulated in response to external stresses caffeine and antifungal agents. Unexpectedly, we find that the N-terminal region of Epe1 is removed in response to insults, resulting in diminished Epe1 heterochromatin nuclear foci, cytoplasmic accumulation and redistribution of heterochromatin to other locations, including those that confer resistance through resulting gene repression. Epe1 cleavage is mediated by E3 ubiquitin ligases and the proteasome in a process resembling regulated <u>u</u>biquitin/proteasome-dependent <u>p</u>rocessing (RUP)^33, 34^. Constitutive tEpe1 expression results in more H3K9me-heterochromatin, increased associated gene repression and elevated resistance. Epe1 processing is dependent on signalling through the cell integrity MAP kinase stress pathway which therefore modulates heterochromatin-mediated caffeine and antifungal resistance. Epe1-related histone demethylase proteins are broadly conserved, suggesting that similar mechanisms contribute to adaption to external insults and antifungal resistance in pathogenic fungi.

## Results

### Epe1 exhibits altered mobility upon exposure to stress

Exposure of fission yeast expressing N-terminally 3xFLAG-tagged Epe1 to caffeine results in reduced levels of this key negative regulator of heterochromatin^13^. However, caffeine treatment of cells expressing C-terminally 13xMyc-tagged Epe1 (Epe1-Myc) revealed a faster-migrating form (hereafter tEpe1) and reduced levels of full-length Epe1-Myc (Fig.1a,b). This effect was independent of the tag as a faster-migrating form of C-terminally GFP-tagged Epe1 (Epe1-GFP) was also detected upon caffeine exposure (Supp. Fig.1a). tEpe1 also appeared upon osmotic (1M KCl) or prolonged oxidative (1mM H2O2) stress (Fig.1b). By comparison, neither amount nor mobility of C-terminally TAP-tagged Lid2 JmjC H3K4 demethylase^35^ were altered by caffeine exposure (Fig.1c). Thus, the mobility of a proportion of Epe1-Myc increases in response to several external stresses.

**Fig. 1:**
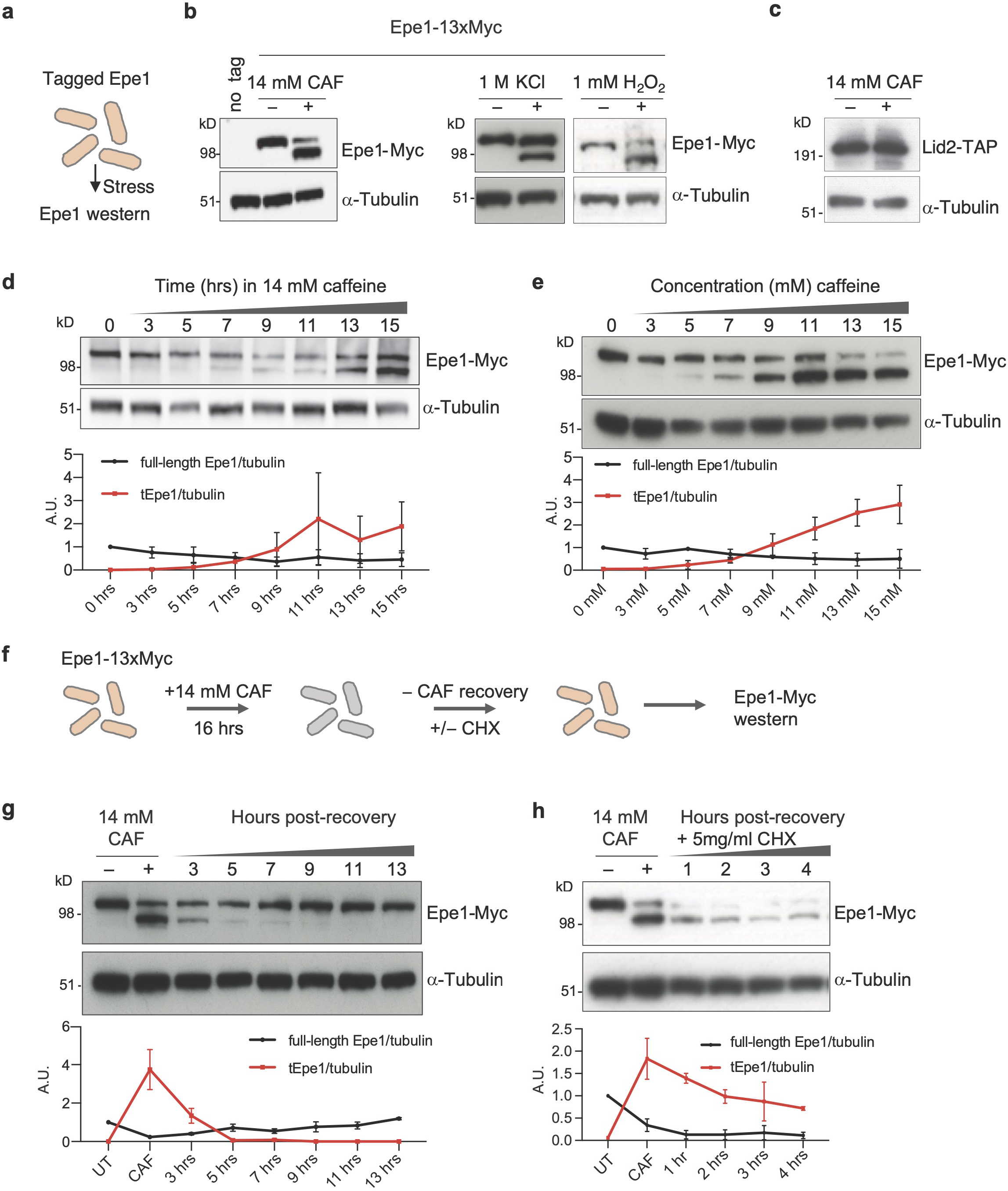
Epe1 migrates faster after caffeine stress and requires protein synthesis for recovery. a. Schematic of cells expressing tagged Epe1 from endogenous promoter exposed to various stresses followed by western. b. C-terminally tagged Epe1-13xMyc or *α*-tubulin western from cells untreated (-)/treated (+) with 14mM caffeine, 1M KCl or 1mM H2O2 for 16h. c. Lid2-TAP or *α*-tubulin western from cells untreated(-)/treated(+) with 14mM caffeine. d. Epe1-13xMyc or *α*-tubulin western from cells incubated for indicated times in 14mM caffeine. e. Epe1-13xMyc or *α*-tubulin western from cells incubated with indicated caffeine concentrations for 16h. f. Cycloheximide block-recovery scheme performed on Epe1-13xMyc cells. g. Epe1-13xMyc or *α*-tubulin western from cells untreated(-)/treated(+) with 14mM caffeine for 16h, following caffeine wash-out and cell recovery in absence of caffeine for indicated times. h. As g. but recovery in 5mg/ml cycloheximide. Plots d,e,g,h: full-length (FL-Epe1) and truncated (tEpe1) levels relative to *α*-tubulin loading control.

The appearance of tEpe1 was time and concentration dependent, with tEpe1 visible after 7h exposure to 14mM caffeine and increasing steadily thereafter, and upon 16h treatment with 5-15mM caffeine (Fig.1d,e). Upon caffeine removal after 14mM/16h treatment, the proportion of tEpe1 declined with time and was undetectable after 9h when levels of normally migrating Epe1-Myc had recovered to those of untreated cells (Fig.1f,g). Cycloheximide prevented recovery, indicating that new protein synthesis is needed to restore Epe1-Myc to the normally migrating form (Fig.1f,h).

### Altered Epe1 mobility results from cleavage and loss of its N-terminal region

Altered protein mobility can result from changes in transcription, translation initiation, post-translational modifications or protein cleavage. *epe1* contains no introns, precluding altered splicing as an explanation (www.pombase.org/gene/SPCC622.16c). RT-qPCR analysis indicated that steady state levels of *epe1* transcript measured either immediately upstream of the initiating AUG or within the coding sequence were unaffected by caffeine (Supp. Fig.1b). Thus, stress-induced tEpe1 is unlikely to result from a downstream transcriptional start site (TSS). Indeed, no change in *epe1*^+^ TSS usage in stress was previously detected^36^.

Mass spectrometry analysis of affinity-selected Epe1-GFP indicated that several residues gained phosphorylation following caffeine treatment (Supp. Fig.1c). However, the mobility of neither normally migrating nor tEpe1 was altered by lambda phosphatase treatment, suggesting that phosphorylation does not cause the stress-mediated increase in Epe1 mobility (Supp. Fig.1d).

To investigate if exposure to caffeine results in post-translational cleavage of Epe1, a strain expressing Epe1 tagged at both the N-terminus (3xMyc) and C-terminus (GFP) from the endogenous locus was constructed (Myc-Epe1-GFP). Anti-GFP westerns following caffeine treatment detected tEpe1-GFP but only the normal, slower-migrating form was detected by anti-Myc western, albeit it at a reduced level (Fig.2a). This observation suggests that in response to stress a region containing the N-terminal 3xMyc tag is cleaved away from Myc- Epe1-GFP to leave both full-length (FL-) Epe1-GFP and truncated tEpe1-GFP detectable.

**Fig. 2:**
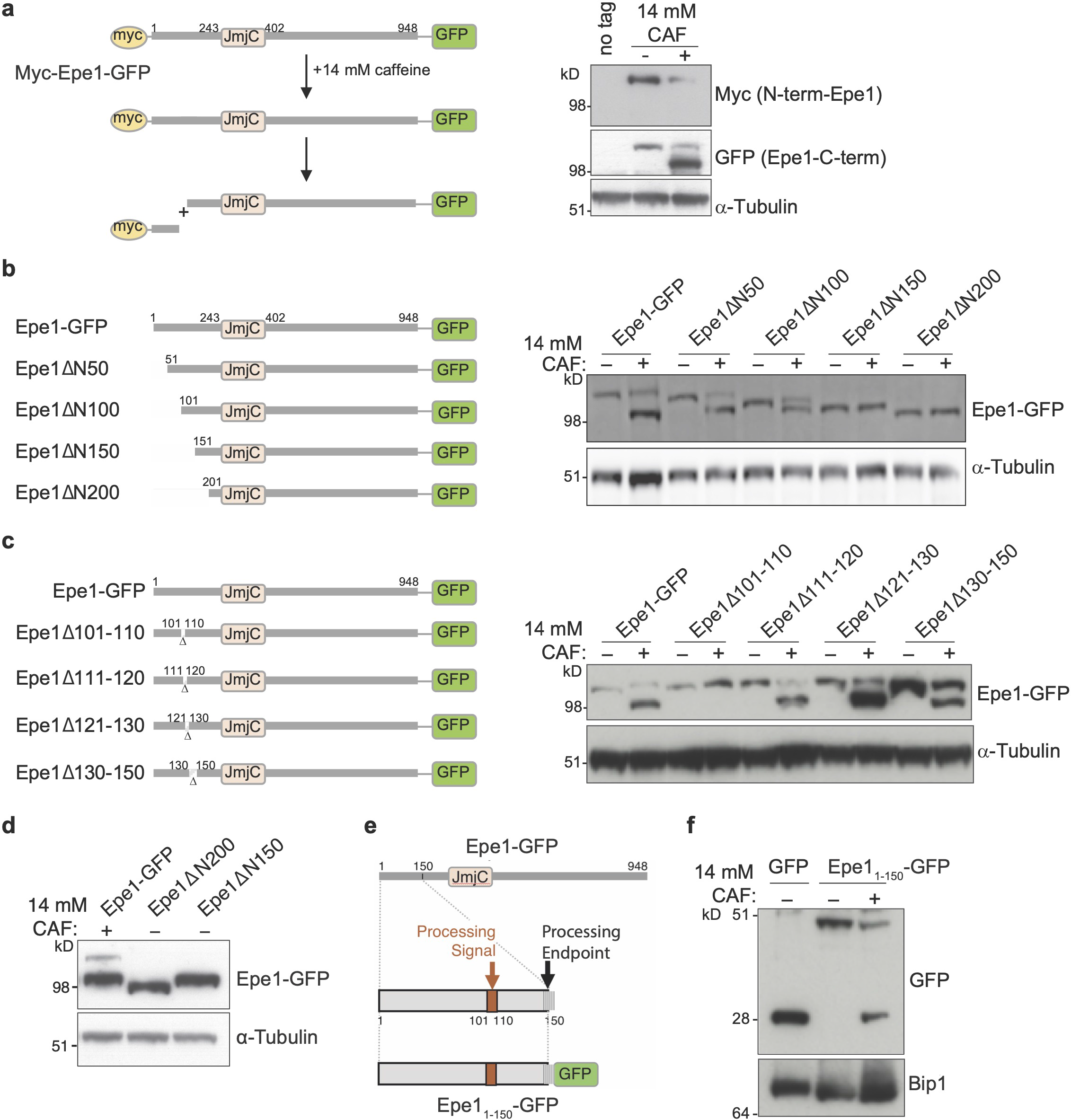
Epe1 altered mobility results from cleavage and release of the N-terminal region. a. Schematic of 3xMyc-Epe1-GFP: cleavage in N-terminal region would release N-terminal 3xMyc from C-terminal GFP-tagged Epe1 of reduced size relative to full length 3xMyc-Epe1- GFP (left). Myc or GFP-tagged 3xMyc-Epe1-GFP, or *α*-tubulin westerns from cells untreated (-)/treated (+) with 14mM caffeine (right). b. Schematic of indicated Epe1 N-terminal truncation mutants expressed as GFP fusions from the endogenous *epe1* locus (left). N-terminal Epe1 truncation-GFP fusion proteins or *α*-tubulin westerns from cells untreated (-)/treated (+) with 14mM caffeine (right). c. Schematic of indicated 10 or 20 residue deletion mutants in the N-terminal coding region of the endogenous *epe1* gene expressed as GFP fusions (left). Western detecting indicated mutant Epe1-GFP fusion proteins or *α*-tubulin from cells untreated (-)/treated (+) with 14mM caffeine (right). d. Western comparing Epe1-GFP migration following 14mM/16h caffeine treatment with constitutively truncated Epe1 and *α*-tubulin loading control. e. Schematic of proposed Epe1 N-terminal processing signal and endpoint. f. Epe1N150-GFP (N-terminal 150 residues of Epe1 fused to GFP) or Bip1 loading control western from cells untreated (-)/treated (+) with 14mM caffeine.

### The N-terminal region of Epe1 carries a portable signal for stress-mediated processing

To investigate the requirements for stress-induced Epe1 cleavage strains expressing a series of N-terminal truncations of Epe1-GFP were generated (Fig.2b). Epe1ΔN50-GFP, Epe1ΔN100-GFP, Epe1ΔN150-GFP, and Epe1ΔN200-GFP truncated proteins were expressed at similar levels to FL-Epe1-GFP. Following caffeine treatment, cells expressing Epe1-GFP, Epe1ΔN50-GFP, or Epe1ΔN100-GFP exhibited faster-migrating forms. However, no caffeine-induced change in Epe1ΔN150-GFP or Epe1ΔN200-GFP mobility was observed (Fig.2b), suggesting that a region required for caffeine-induced cleavage resides between Epe1 residues 100 and 150.

Further strains, expressing Epe1-GFP proteins lacking 10 or 20 residues between residues 100 and 210 (Fig.2c; Supp. Fig.2a) were subjected to western analysis following caffeine treatment. Removal of residues 101-110 prevented stress-mediated cleavage of Epe1Δ101- 110-GFP whereas FL-Epe1-GFP, Epe1Δ111-120-GFP, Epe1Δ111-130-GFP and Epe1Δ130-150-GFP, Epe1Δ151-170-GFP, Epe1Δ171-190-GFP and Epe1Δ190-210-GFP all exhibited a faster-migrating form. These analyses suggest that residues 101-110 are required for Epe1 cleavage. Direct comparison of caffeine-induced tEpe1 migration with Epe1ΔN150-GFP and Epe1ΔN200-GFP indicated that, although residues 101-110 are necessary for Epe1 processing, the new N-terminus resides near residue 150 (Fig.2d,e). A precise cleavage site in Epe1 could not be defined by mass spectrometry. However, after Epe1-Myc immunoprecipitation (three replicates), three peptides (residues: 52-57, 57-67, 83-96) from within the first 150 amino acids of FL-Epe1-Myc were identified in cells grown without caffeine with high confidence (1% FDR) but were not detectable from tEpe1 following caffeine treatment. In contrast, fifteen peptides beyond residue 150 were detected with high confidence (1% FDR) from both FL-Epe1-Myc and tEpe1-Myc (Supp. Fig.2b).

To determine if the N-terminal region is sufficient to mediate caffeine-induced cleavage the first 150 residues of Epe1 were fused to GFP (Epe11-150-GFP; Fig.2e). Treatment with caffeine released an entity from Epe11-150-GFP that co-migrated with recombinant GFP (27kDa, Fig.2f). We conclude that the first 150 residues of Epe1 are necessary and sufficient to drive cleavage of a heterologous protein in response to external stress.

### Caffeine-induced Epe1 N-terminal removal is proteasome dependent

To gain insight into the mechanism of Epe1 stress-mediated cleavage we affinity selected Epe1-GFP from cells grown with or without 14mM caffeine and applied proteomics to identify associated proteins. Notably, caffeine treatment resulted in increased association of 23 of 35 known proteasome subunits^38^, representing components of both the 20S core (9) and 19S regulatory (14) particles, with Epe1-GFP (Fig.3a). Caffeine-induced cleavage of 3xMyc-Epe1- GFP should release an N-terminal product of approximately 180 residues with a predicted mass of ∼20kDa (including 3xMyc). However, no such product was detected, suggesting that the N-terminal fragment is rapidly degraded by the proteasome following its initial cleavage (Supp. Fig.3a). To test proteasome involvement in stress-mediated post-translational processing of Epe1 we examined Epe1-13xMyc processing in cells harbouring the *mts2-1* mutation in the Rpt2/ATPase subunit of the 19S proteasome regulatory particle^39^. tEpe1 was generated in wild-type but not *mts2-1* cells grown at the permissive temperature, indicating that proteasome function contributes to caffeine-dependent Epe1 cleavage (Fig.3b). A shorter Mst2 histone acetyltransferase (HAT) isoform is made in response to stress by a distinct mechanism involving use of a downstream TSS^13, 36^. Production of N-terminally truncated Mst2-13xMyc in caffeine was unaffected by the *mts2-1* mutant: indicating that proteasome-mediated processing is Epe1 specific and that *mts2-1* does not prevent other changes in response to caffeine stress (Supp. Fig.3b).

**Fig. 3:**
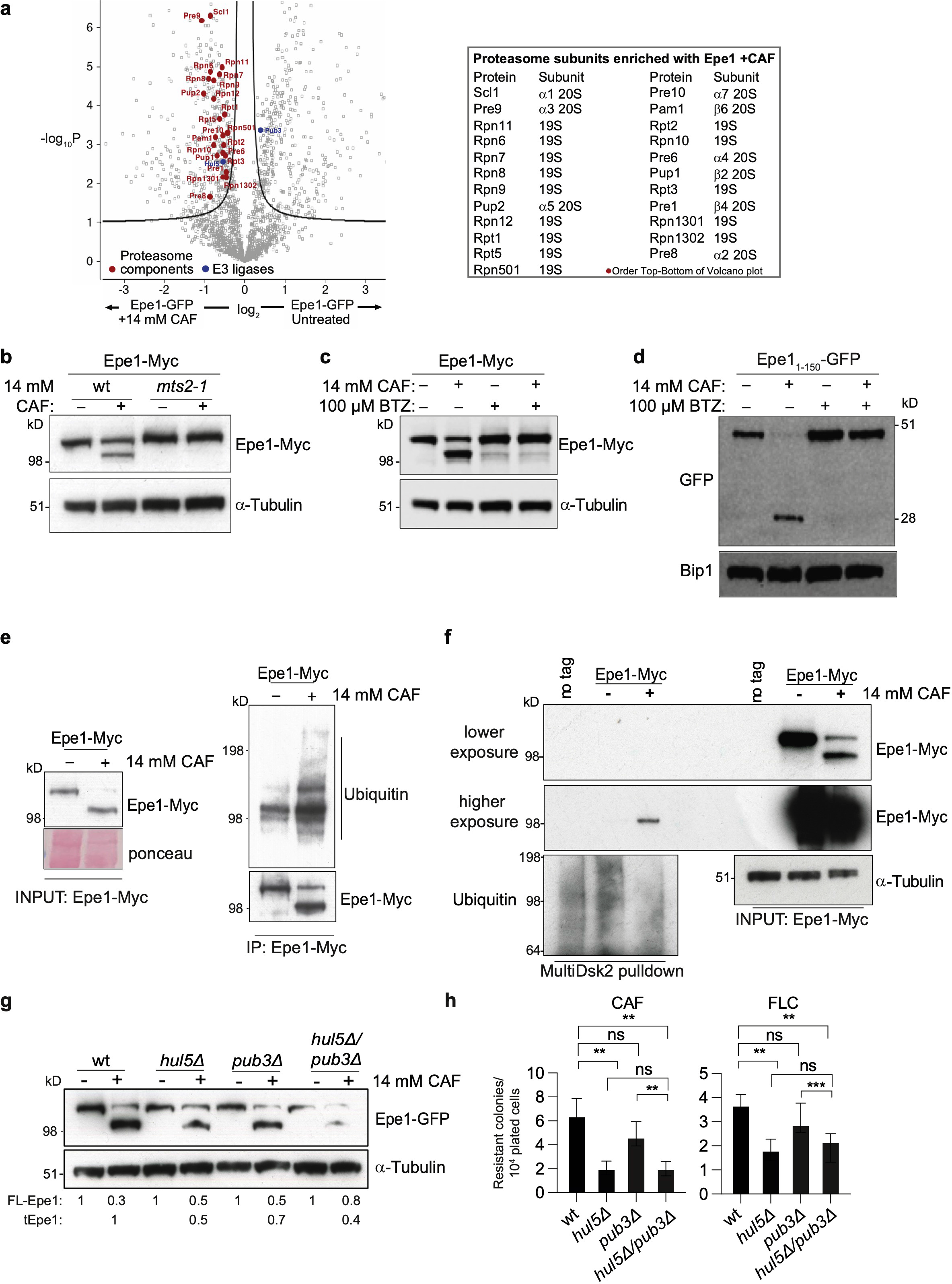
The Epe1 N-terminal region is sufficient for caffeine-induced cleavage that is dependent on proteasome function. a. Volcano plot showing proteins enriched with Epe1-GFP by proteomic analysis of extracts from cells untreated or treated with 14mM/16h caffeine. Red dots: 26S proteasome components enriched with Epe1-GFP after caffeine treatment named in table (right). Blue dots: enriched E3 ubiquitin ligases. Full data, Supp. Table 3. b. Epe1-GFP western from wild-type (wt) or proteasome defective cells (*mts2-1*) untreated (-)/treated (+) with 14mM caffeine. c. Epe1-Myc western from cells untreated (-)/treated (+) with 14mM caffeine and/or proteasome inhibitor bortezomib (BTZ). d. Western detecting Epe11-150-GFP following treatments as in c, or recombinant GFP. e. Western of immunoprecipitated Epe1-13xMyc probed with anti-Myc or anti-ubiquitin antibodies in absence (-)/presence(+) of 14mM caffeine. f. Western of Multidsk2 affinity resin enriched proteins probed with anti-Myc or anti-ubiquitin antibodies in absence (-)/presence(+) of 14mM caffeine. g. Western detecting Epe1-GFP or *α*-tubulin in wild-type, *hul5Δ, pub3Δ* or *hul5Δ pub3Δ* cells untreated (-)/treated (+) with 14mM caffeine for 16h. h. Number of resistant colonies formed/1x10^4^ viable cells by wild-type, *hul5Δ, pub3Δ* or *hul5Δ pub3Δ* cells plated on caffeine or fluconazole plates. Statistical analysis of three biological replicates performed as in Methods.

The proteasome inhibitor bortezomib primarily inhibits the Pts1/*β*5 peptidase of the 20S core particle^40^. Addition of bortezomib prevented processing of Epe1-Myc to tEpe1 in the presence of caffeine, and also inhibited caffeine-induced release of GFP from Epe11-150-GFP (Fig.3c,d). In contrast, bortezomib did not inhibit caffeine-induced Mst2/HAT truncation (Supp. Fig.3c), indicating that stress-induced protein truncation is not generally inhibited by defective proteasome function.

Epe1 is a nuclear protein concentrated in 2-4 heterochromatic foci representing clustered centromeres, telomeres and the silent mating type locus^28, 41^ . Treatment of wild-type cells with caffeine resulted in loss of all major nuclear Epe1-GFP foci but not the distinct centromere-specific histone CENP-A^Cnp1^ signal (Supp. Fig.3d,e). Nuclear Epe1-GFP accumulated in proteasome compromised *mts2-1* cells (permissive temperature), consistent with the known Cul4-Ddb1^Cdt2^-mediated ubiquitin-dependent removal of Epe1 from within major blocks of heterochromatin^42^. Although nuclear Epe1 levels were somewhat decreased by addition of caffeine to *mts2-1* cells, substantial amounts remained. Thus, proteasome function, and presumably proteasome-dependent Epe1 processing, is required to deplete nuclear Epe1 in response to caffeine stress.

Following caffeine treatment increased ubiquitin associated with affinity-purified Epe1-Myc and Epe1-Myc was enriched on ubiquitin-binding beads (Fig.3e,f). Thus, ubiquitylation of Epe1 is associated with Epe1-to-tEpe1 caffeine-induced proteasome-dependent processing. The Cul4-Ddb1^Cdt2^ E3 ubiquitin ligase targets Epe1 for degradation within major heterochromatin domains^42^, however, Ddb1^Cdt2^ loss had little impact on Epe1-to-tEpe1 processing (Supp. Fig.3f) suggesting involvement of other E3 ligases. Our proteomics analysis of affinity-selected Epe1-GFP detected enrichment of the Hul5 and Pub3 E3 ubiquitin ligases in the presence or absence of caffeine, respectively (Fig.3a). Cells lacking Hul5 exhibited decreased caffeine-induced Epe1-to-tEpe1 processing, whereas deletion of *pub3^+^* had little impact alone or in combination with *hul5*Δ. Thus Hul5, but not Pub3, E3 ubiquitin ligase contributes to proteasome-mediated Epe1 processing in response to caffeine (Fig.3g). Consistent with decreased Epe1-to-tEpe1 processing in response to external stress, both *hul5*Δ and *hul5*Δ *pub3*Δ cells, but not *pub3*Δ or *ddb1*Δ cells, exhibited a lower frequency of caffeine and fluconazole resistant colonies compared to wild-type cells (Fig.3h). We conclude that Hul5 E3 ubiquitin ligase contributes to the removal of the N-terminal region from Epe1 to generate tEpe1 by promoting its ubiquitin-dependent proteasome-mediated processing.

### N-terminally truncated Epe1 accumulates in the cytoplasm

To further investigate the cellular location of processed Epe1-GFP we utilized constitutively expressed N-terminal truncations. Epe1ΔN100-GFP gave a similar pattern and nuclear signal intensity to FL-Epe1-GFP. Removal of an additional 50 (Epe1ΔN150-GFP) or 100 (Epe1ΔN200-GFP) residues resulted in complete loss of nuclear GFP foci, reduced nuclear signal intensity, with a corresponding increase in cytoplasmic signal intensity (Fig.4a,b). These analyses indicate that the N-terminal 150 residues of Epe1 are required to maintain normal nuclear Epe1 levels and its concentration at heterochromatin domains, and that their removal increases the cytoplasmic location of tEpe1.

**Fig. 4:**
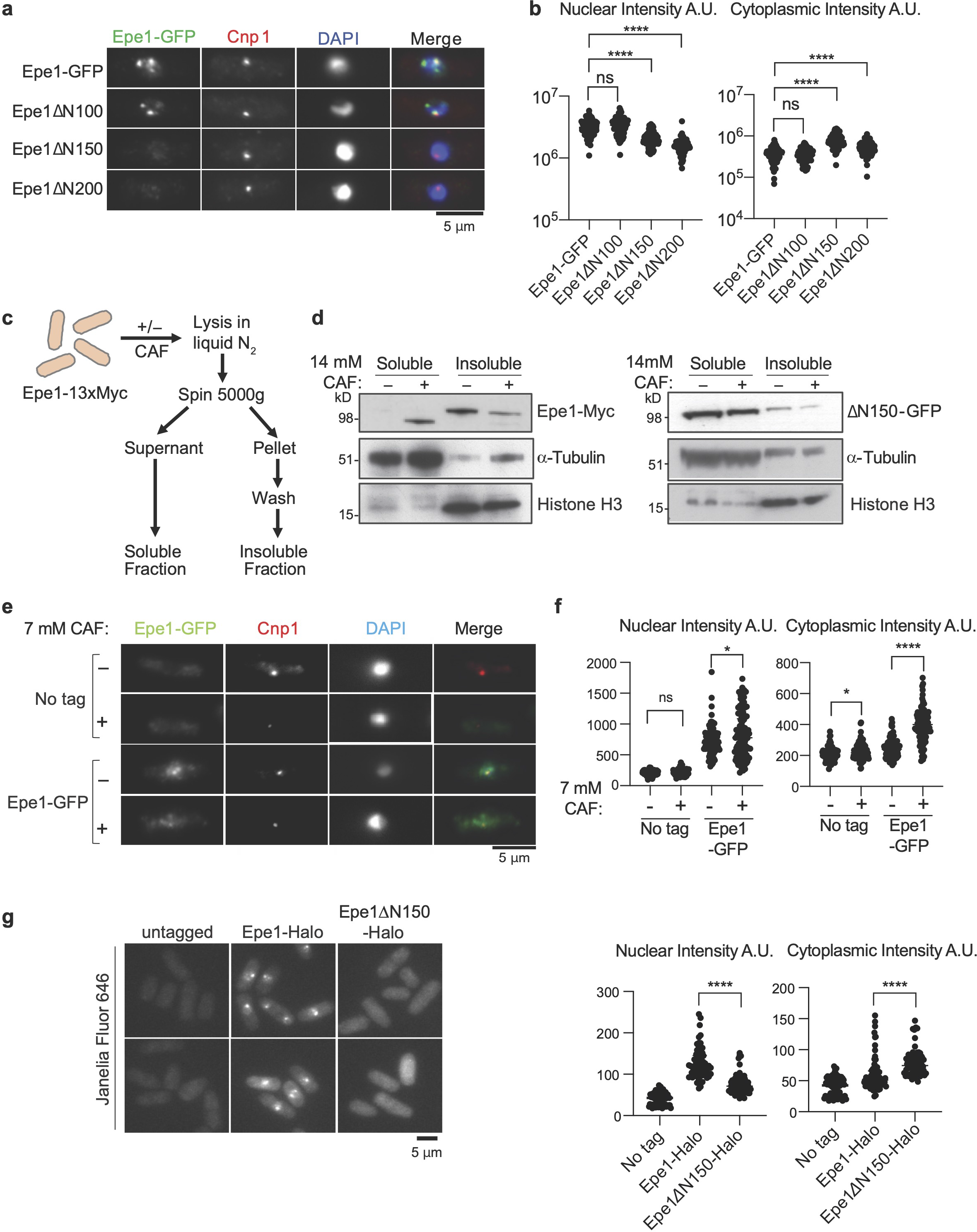
N-terminally truncated Epe1 and caffeine-processed Epe1 lose association with chromatin. a. Immunolocalization of Epe1-GFP, indicated N-terminal Epe1 truncation-GFP fusion proteins (green) and CENP-A^Cnp1^ control (red) in DAPI stained (blue) cells. Scale bar, 5*μ*m. b. Quantification of anti-GFP nuclear and cytoplasmic signals for cells from a. 100 cells analysed per sample. c. Schematic of insoluble chromatin and soluble cytoplasmic fractionation. d. Epe1-13xMyc (left) or Epe1ΔN150-GFP (right), histone H3 or *α*-tubulin westerns on soluble and insoluble fractions from cells untreated (-)/treated (+) with 14mM caffeine. e. Immunolocalization as in a. of Epe1-GFP in cells untreated (-)/treated (+) with 7mM caffeine compared with control untagged cells. f. Quantification of anti-GFP nuclear or cytoplasmic signals for cells from e. 100 cells analysed per sample. g. Images of untagged, Epe1-Halo and Epe1ΔN150-Halo cells exposed to the JF646 fluorescent dye (left) and quantification of nuclear or cytoplasmic signals (right). 100 cells analysed/sample. Statistical analysis of three biological replicates performed as in Methods.

Lysates from cells expressing Epe1-Myc grown with or without caffeine were fractionated into soluble cytosolic supernatants and insoluble chromatin-containing pellets (Fig.4c,d). As expected, most *α*-tubulin was released into the supernatant whereas chromatin (histone H3) was retained in the pellet. Without caffeine most Epe1 stayed in the insoluble pellet, as expected for this chromatin-associated protein. Upon caffeine-induced processing, most tEpe1 was present in the soluble fraction whereas unprocessed Epe1-Myc remained in the pellet. This finding indicates that caffeine-induced removal of Epe1’s N-terminus results in its loss from chromatin. In agreement with this observation, constitutively truncated Epe1ΔN150- GFP was detected mainly in the soluble fraction and was unaffected by caffeine treatment. Moreover, comparison of full-length Epe1-GFP and Epe1ΔN150-GFP-associated proteomes revealed that they occupy distinct cellular compartments (Supp. Fig.4a,b). Epe1-GFP associated with several other chromatin-associated nuclear proteins, including histones, the chromodomain protein Swi6, and the bromodomain proteins Bdc1 and Bdf2. In contrast, constitutively truncated Epe1ΔN150-GFP (equivalent to the caffeine-induced tEpe1-GFP) was associated with cytoplasmic proteins, consistent with a more cytosolic location.

To minimize the deleterious effects of caffeine-induced stress on cell morphology and cytology, the location and cytoplasmic/nuclear signal intensity of Epe1-GFP was assessed in the presence and absence of a lower caffeine concentration (7mM). Under these less stressful conditions less Epe1-to-tEpe1 processing occurs than at higher caffeine concentrations (Fig.1e), and most Epe1-GFP remained in the nucleus, nevertheless, cytoplasmic Epe1-GFP signals were elevated (Fig.4e,f). Moreover, employment of the significantly brighter Halo-tag in place of GFP confirmed FL-Epe1-Halo to be primarily nuclear whereas Epe1ΔN150-Halo was distributed throughout the nucleus and cytoplasm (Fig.4g). Together these data indicate that stress-mediated Epe1-to-tEpe1 processing, for which Epe1ΔN150-GFP and Epe1ΔN150- Halo provide constitutively expressed proxies, results in loss of tEpe1 from chromatin and its increased presence in the cytoplasm.

### Removal of the Epe1 N-terminal region increases heterochromatin formation

Exposure of fission yeast to caffeine reduces Epe1 association with constitutive heterochromatin regions, alters the distribution of H3K9me-dependent heterochromatin and instigates caffeine resistance through the repression of underlying genes^13^. We next investigated if caffeine-induced N-terminal Epe1 truncation results in changes to its chromatin distribution and that of heterochromatin. Silencing assays at the euchromatin-heterochromatin border on the left side of *cen1* revealed that, as in *epe1*Δ cells, repression of the nearby inserted euchromatic *ura4^+^* marker gene occurred in cells expressing Epe1ΔN150 (growth on counter-selective FOA plates; Supp. Fig.5a). Quantitative ChIP showed that Epe1-GFP was enriched at centromere repeats (*dg*), a *cen1* adjacent boundary region (*IRC1-L*) and a region near a telomere (*tlh2*), where Epe1 normally associates^22, 42^ (Fig.5a). In contrast, removal of Epe1 N-terminal 150 residues (Epe1ΔN150-GFP) resulted in loss of association with these regions (Fig 5a). As expected, *epe1Δ* cells exhibit higher H3K9me2 levels at known facultative heterochromatin islands^22^ (*isl3*, *isl6*; Fig.5b). However, cells expressing only Epe1ΔN150-GFP exhibited greater H3K9me2 levels at several islands (*isl1, isl3, isl6, isl9*). Importantly, cells expressing non-cleavable Epe1ΔN101-110-GFP exhibited no change in its association with major heterochromatin regions compared to Epe1-GFP before or after caffeine treatment (*dg, IRC, tel2R*; Fig.5a,c) nor in H3K9me2 levels at *dg* centromere repeats or any island examined (Fig.5b). Thus removal of the Epe1 N-terminal region mimics caffeine treatment^13^ and Epe1 loss^22^ in increasing H3K9 methylation-dependent heterochromatin at various chromosomal locations.

**Fig. 5:**
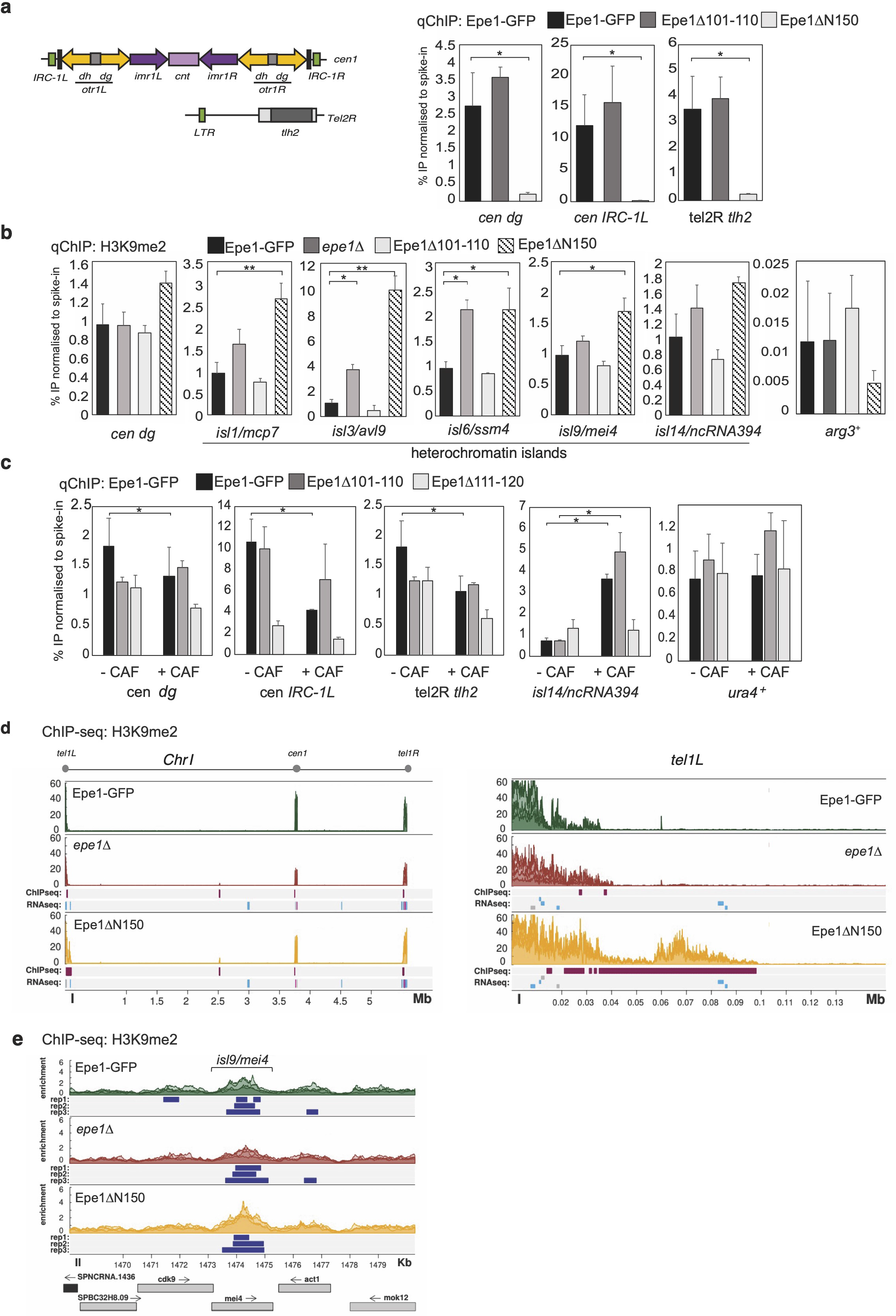
Loss of Epe1 nuclear foci coincides with reduced heterochromatin association, increased H3K9 methylation and associated gene expression changes. a. Diagram of centromere and telomere heterochromatin regions (left). ChIP-qPCR for Epe1- GFP, Epe1ΔN100-GFP, Epe1ΔN101-110-GFP or Epe1ΔN150-GFP levels at centromeric outer repeats (*dg*), a boundary region (*IRC1-L*) and telomere (*tlh2*). Epe1-GFP protein levels were normalised to *S. cerevisiae* Sgo1-GFP spike-in control. b. ChIP-qPCR for H3K9me2 levels at *dg* centromere repeats, known heterochromatin islands (*isl1, isl3, isl6, isl9, isl14*) and euchromatic gene (*arg3^+^*) in cells with the indicated manipulations at *epe1*. H3K9me2 levels were normalised to an *S. octosporus* spike-in control. c. ChIP-qPCR for Epe1-GFP, Epe1ΔN101-110-GFP or Epe1ΔN111-120-GFP levels at *dg* centromere repeats, *cen1* boundary region (*IRC1-L*), telomere adjacent tlh2, *isl14/ncRNA.394*, and a euchromatic gene (*ura4^+^*). d. ChIP-seq showing H3K9me2 distribution in Epe1-GFP, *epe1Δ,* and Epe1ΔN150-GFP cells across chromosome 1 (*Chr1*, left) and adjacent to the left telomere (*tel1L*, right). Three colour shades represent each biological replicate. H3K9me2 ChIP-seq or RNA-seq reads in *epe1Δ* or Epe1ΔN150-GFP cells relative to wild-type Epe1-GFP are shown in tracks below. Features, genomic bins (ChIP-seq) or annotated genes (RNA-seq) with significantly (pvalue <0.1; Log2 fold change >2) increased (magenta) or decreased (cyan) read counts are indicated. e. Three ChIP-seq biological replicates show variable expansion of H3K9me2 over island *isl9/mei4* locus of *epe1*Δ and Epe1ΔN150-GFP cells relative to wild-type Epe1-GFP. MACS2 detected peaks with enrichments >2 are shown (blue). For ChIP-qPCR a-c: Mean, 3 biological replicates; error bars, standard deviations. Significance of the difference between samples was evaluated using Student’s *t*-test. (*) *P* <0.0.33, (**) *P* <0.002; (***) *P* <0.0002; (****) *P* <0.0001; (n.s). not significant.

ChIP-seq analysis confirmed that H3K9me2 levels increase at sub-telomeric and island regions in *epe1Δ* null and Epe1ΔN150-GFP expressing cells (Fig.5d,e). Consistent with ChIP-qPCR analysis (Fig. 5b), ChIP-seq also shows that at some locations heterochromatin spreads further in Epe1ΔN150-GFP cells relative to *epe1Δ* cells, indicating that Epe1ΔN150- GFP somehow promotes increased heterochromatin formation (*tel1L, isl9/mei4*; Fig.5d,e). This increased spreading is particularly prominent at sub-telomeric regions and RNA-seq shows that many telomere proximal genes display even greater repression in Epe1ΔN150- GFP cells compared to *epe1Δ* cells (Supp. Figure 5b,c). Therefore, since Epe1ΔN150-GFP cells are not equivalent to *epe1Δ* cells, these data reveal that Epe1ΔN150-GFP, a surrogate for tEpe1, retains functions that can regulate heterochromatin distribution.

### Truncated Epe1 and associated catalytic activity enhance caffeine and antifungal resistance through increased heterochromatin formation

To further explore the influence of Epe1 processing on resistance of fission yeast to caffeine and antifungals we measured the frequency of caffeine resistance amongst viable cells expressing truncated or mutant forms of Epe1. Like *epe1*Δ, cells expressing Epe1ΔN150-GFP displayed an increased frequency of caffeine-resistant colonies relative to cells expressing wild-type Epe1-GFP or Epe1ΔN100-GFP (Fig.6a). In contrast, cells expressing non-cleavable Epe1Δ101-110-GFP, which in response to caffeine remains enriched at major heterochromatic regions (dg, IRC-1L, tel2R; Fig.5c), exhibited greater sensitivity to caffeine than wild-type cells (Fig.6b). Removal of Clr4 H3K9 methyltransferase from *epe1*Δ, Epe1ΔN100-GFP or Epe1ΔN150-GFP cells reduced the frequency of caffeine-resistant colonies, demonstrating that heterochromatin is required to confer resistance^13^ (Fig.6a). Thus, Epe1 N-terminal truncation to residue 150 results in its increased cytoplasmic location and phenocopies the H3K9me heterochromatin-dependent resistance phenotype of *epe1Δ* cells. Reciprocally, non-cleavable Epe1Δ101-110-GFP remains chromatin associated in the presence of caffeine, thus the reprogramming required to elicit resistance cannot take place, rendering these cells super-sensitive.

**Fig. 6.**
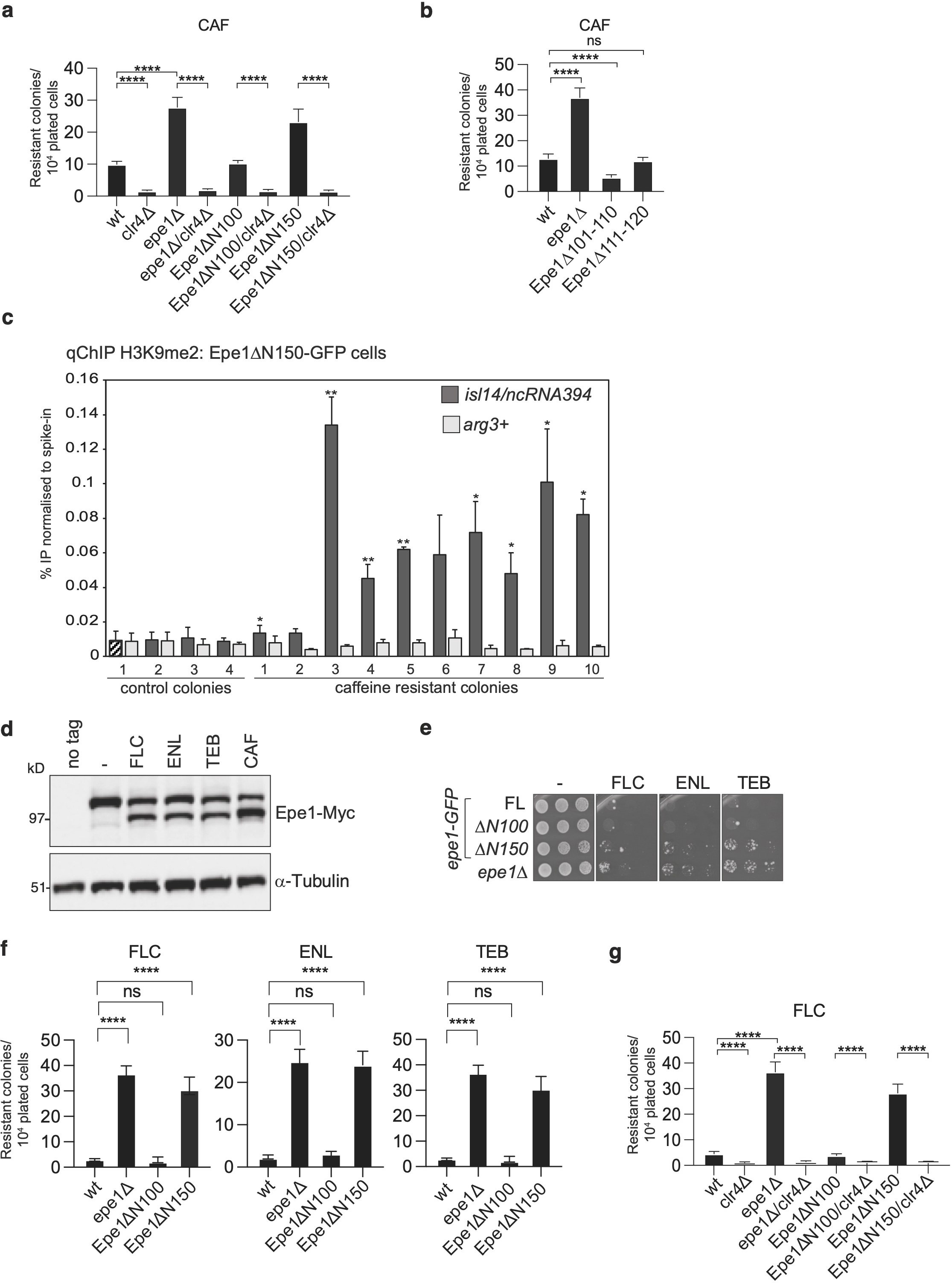
Clr4-dependent H3K9 methylation mediates caffeine and antifungal resistance, and is enhanced by Epe1 truncation. a. Number of resistant colonies formed/1x10^4^ viable cells plated by wild-type, *epe1Δ, epe1Δ/clr4Δ,* Epe1ΔN100-GFP, Epe1ΔN50-GFP*/clr4Δ*, Epe1ΔN150-GFP and Epe1ΔN150- GFP/*clr4*Δ on plates containing 16mM caffeine (CAF). b. Number of resistant colonies formed/1x10^4^ viable cells plated by wild-type, *epe1Δ* Epe1Δ101-110-GFP and Epe1Δ111-120-GFP cells on plates containing 16mM caffeine (CAF). c. ChIP-qPCR for H3K9me2 at the island 14/ncRNA394 or *arg3^+^* loci on ten caffeine-resistant and four control colonies (from no caffeine) formed by Epe1ΔN150-GFP cells. Signals were normalised to an *S. octosporus* spike-in control and significance calculated to control colony 1 (cross-hatched). Mean, 3 biological replicates; error bars indicate standard deviation. d. Epe1-13xMyc or *α*-tubulin westerns from cells untreated (-)/treated (+) with 0.5mM fluconazole (FLC), 6*μ*M enilconazole (ENL), 16*μ*M tebuconazole (TEB) or 14mM CAF e. Serial dilution growth assay of cells with indicated manipulation at *epe1* on plates containing no antifungal (-), FLC, ENL, or TEB. f. Number of resistant colonies formed/1x10^4^ viable cells with indicated manipulation at *epe1* on plates containing FLC, ENL or TEB. g. Number of resistant colonies formed/1x10^4^ viable cells plated by wild-type, *clr4Δ, epe1Δ epe1Δ/clr4Δ,* Epe1ΔN100-GFP, Epe1ΔN50-GFP*/clr4Δ*, Epe1ΔN150-GFP and Epe1ΔN150- GFP*/clr4Δ* on plates containing 0.5mM fluconazole (FLC).

ChIP-qPCR performed on caffeine-resistant isolates from Epe1ΔN150-GFP cells revealed significantly higher H3K9me2 levels over the *isl14/ncRNA394* locus in eight of ten isolates analyzed (Fig.6c). Previously we showed that caffeine resistance directly results from heterochromatin-associated gene repression over *isl14/ncRNA394*^13^. We therefore conclude that cells expressing constitutively truncated Epe1ΔN150-GFP exhibit increased caffeine resistance due to similar heterochromatin-mediated epimutations.

Caffeine resistant epimutants were previously shown to also exhibit resistance to clinical and agricultural crop-plant antifungals^13^. As with caffeine and salt stress (Fig.1b), treatment with a clinical antifungal (fluconazole, FLC), or crop-plant fungicides enilconazole (ENL) or tebuconazole (TEB), induced Epe1-to-tEpe1 processing (Fig.6d). Moreover, N-terminally truncated Epe1ΔN150-GFP, but not Epe1ΔN100-GFP, increased the frequency of colonies resistant to these antifungals to levels similar to *epe1Δ* cells (Fig.6e,f). Removal of Clr4 from *epe1*Δ, Epe1ΔN100-GFP or Epe1ΔN150-GFP cells reduced the frequency of fluconazole-resistant colonies, indicating that this increase in anti-fungal resistance also requires H3K9me- dependent heterochromatin (Fig.6g).

The phenotype of cells expressing constitutively truncated Epe1ΔN150-GFP appears similar to cells completely lacking Epe1 (*epe1*Δ) in that heterochromatin accumulates at particular chromosomal locations and both exhibit a Clr4-dependent increase in caffeine and antifungal resistance. However, insult-induced tEpe1 and Epe1ΔN150 retain an intact JmjC demethylase domain and accumulate in the cytoplasm (Fig.4). Moreover, cells expressing Epe1ΔN150- GFP exhibit elevated H3K9me2 levels at certain locations (Fig.5b,d). To determine if the JmjC domain of Epe1ΔN150 contributes to their resistance we constructed cells expressing a predicted catalytically dead version of FL-Epe1-GFP and Epe1ΔN150-GFP bearing H297A and K314A substitutions to disrupt iron/Fe(II) and 2-oxoglutarate binding, respectively (Epe1- cd-GFP, Epe1ΔN150-cd-GFP). Both Epe1-cd-GFP and Epe1ΔN150-cd-GFP were expressed from the *epe1* locus at similar levels to their catalytically active counterparts (Supp. Fig.6a). However, cells expressing Epe1-cd-GFP (naturally induced tEpe1) or Epe1ΔN150-cd-GFP (constitutive tEpe1) displayed reduced frequencies of caffeine and fluconazole resistant colonies relative to Epe1-GFP and Epe1ΔN150-GFP cells (Supp. Fig.6b). Since tEpe1/Epe1ΔN150 are more cytoplasmically localized, their intact JmjC demethylase domains may promote heterochromatin-mediated resistance via other, perhaps cytoplasmic, substrates. Detailed future investigation will be required to identify potential additional Epe1 substrates and how they might contribute to resistance phenotypes.

### The cell integrity MAP kinase pathway regulates stress-induced proteasome-mediated Epe1 cleavage

TOR and MAP kinase-dependent signalling pathways sense and mediate adaptation of fission yeast to the external environment. TOR serine/threonine kinase dependent pathways are major regulators of cellular metabolism, growth and cell cycle in response to nutrient availability and other stresses^43, 44^. TORC2 signalling acts through the single downstream Gad8 effector kinase which is chromatin-associated and known to influence heterochromatin integrity^45, 46^ (Supp. Fig.7a). However, *gad8Δ* cells exhibited relatively normal Epe1-to-tEpe1 processing but a reduced frequency of caffeine resistant colonies, suggesting that TORC2- mediated signalling may counteract the effects Epe1 processing has on heterochromatin (Supp. Fig.7b,d). Indeed, TORC2/Gad8 is known to antagonise Epe1 function by promoting heterochromatin formation^45^.

Two fission yeast MAP kinase signalling pathways elicit responses to extracellular changes, resulting in altered gene expression enabling cell adaptation and survival^47–49^. The general stress-activated (SAP, Fig.7a) and the cell integrity (CIP, Fig.7b) pathways are interconnected, and signal through Sty1 and Pmk1 MAP kinases, respectively, to activate the key downstream transcription factor Atf1. Loss of Pek1/MAPKK or Pmk1/MAPK, but not Sty1/MAPK, dramatically reduced tEpe1-GFP generation in the presence of caffeine, retaining FL-Epe1- GFP levels similar to wild-type (Fig.7c,d). Cells lacking Atf1 also retained high levels of FL- Epe1, although some cleavage to tEpe1 was apparent. In contrast, cells lacking the Pmp1 phosphatase, which inhibits Pmk1, displayed even lower levels of FL-Epe1 than wild-type cells because more caffeine-induced tEpe1 was evident (Fig.7d). These results indicate that the CIP pathway is mainly responsible for regulating Epe1 processing in response to external caffeine. Epe1-to-tEpe1 processing in response to fluconazole also required intact CIP/MAPK signalling but in this case Atf1 was required for tEpe1 generation (Supp. Fig.7c); cells lacking Atf1 displayed an equivalent frequency of caffeine-resistant colonies but higher frequency of fluconazole-resistant colonies, compared to wild-type cells (Supp. Fig.7d,e). These observations suggest that caffeine and fluconazole activate Epe1 cleavage through overlapping routes.

**Fig. 7:**
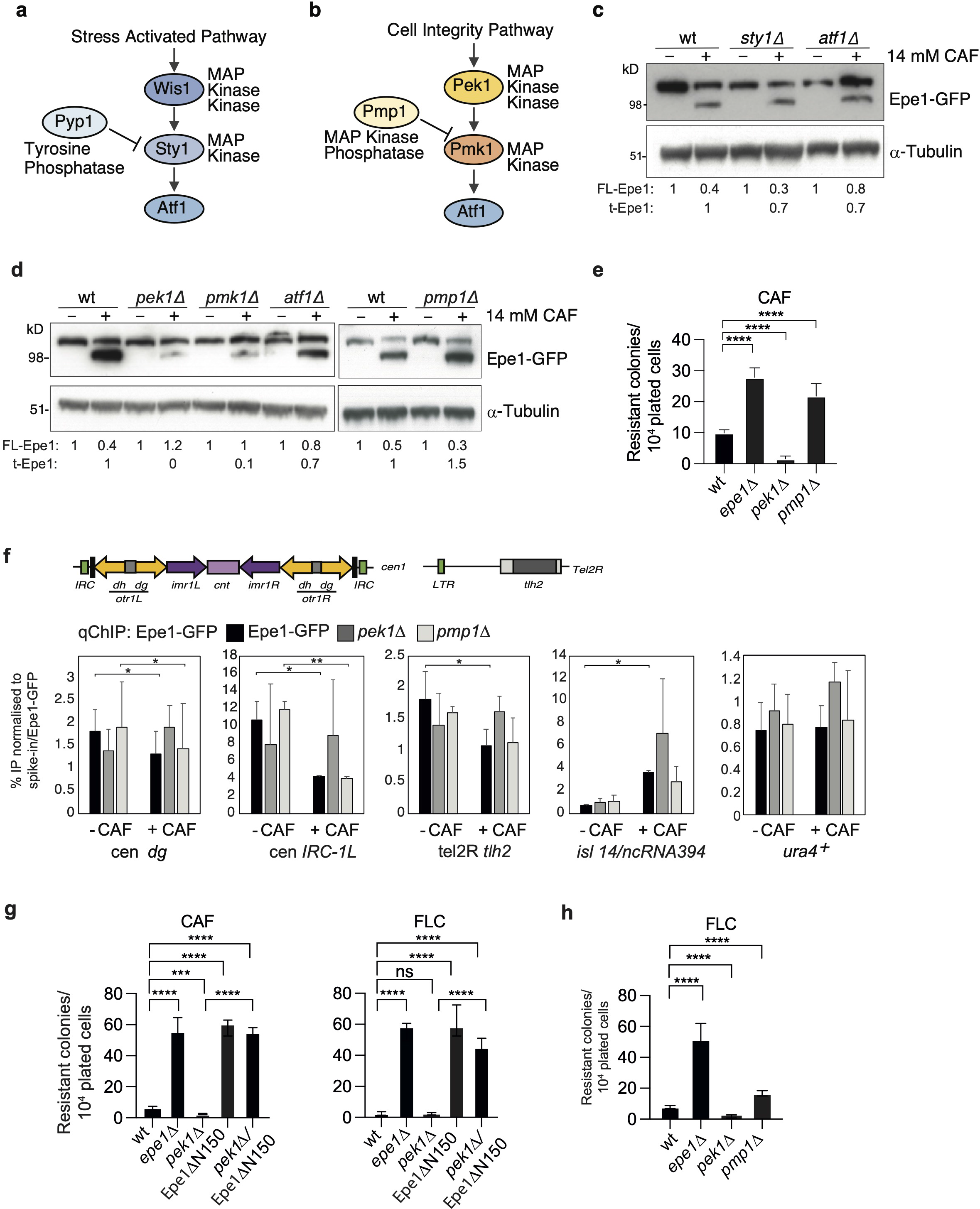
The cell integrity stress pathway regulates Epe1 processing via Pek1 MAPKK, Pmk1 MAPK and Pmp1 MAPK phosphatase. a. Diagram of the *S. pombe* stress-activated signalling pathway (SAP) end point. b. Diagram of *S. pombe* cell integrity signalling pathway (CIP) end point. c. Epe1-GFP or *α*-tubulin western from wild-type, *sty1Δ,* or *atf1Δ* cells untreated (-)/treated (+) with 14mM caffeine. d. Epe1-GFP or *α*-tubulin western from wild-type, *pek1Δ*, *pmk1Δ, atf1Δ* or *pmp1Δ* cells untreated (-)/treated(+) with 14mM caffeine. e. Number of resistant colonies formed/1x10^4^ viable cells by wild-type, *epe1Δ, pek1Δ* and *pmp1Δ* cells on caffeine plates. f. ChIP-qPCR for Epe1-GFP at the indicated locations in wild-type, *pek1Δ*, or *pmp1Δ* cells cells untreated (-)/treated (+) with 14 mM caffeine. Epe1-GFP levels normalised to *S.cerevisiae* Sgo1-GFP spike-in control. Mean, 3 biological replicates; error bars, standard deviations. g. Number of resistant colonies formed/1x10^4^ viable cells by wild-type, *epe1Δ, pek1Δ*, Epe1ΔN150 and *pek1Δ* Epe1ΔN150 cells plated on caffeine or fluconazole plates. h. Number of resistant colonies formed/1x10^4^ viable cells plated by wild-type, *epe1Δ, pek1Δ* and *pmp1Δ* on plates containing 0.5mM fluconazole (FLC). Statistical analysis of three biological replicates performed as in Methods.

Consistent with Pmk1 and Pmp1 promoting and hindering stress-induced Epe1 cleavage respectively, Epe1-GFP association with major heterochromatic regions in response to caffeine decreased similarly in wild-type and *pmp1Δ* cells (more cleavage), but not *pek1Δ* cells (less cleavage) (*cen dg, IRC1-L*; Fig.7f). Moreover, as in *epe1Δ* cells, *pmp1Δ* cells exhibited an increased frequency of caffeine and fluconazole resistant colonies compared to wild-type (Fig.7e,h), consistent with increased Epe1-to-tEpe1 processing (Fig.7d; Supp. Fig.7c). Reciprocally, loss of the upstream Pek1/MAPKK resulted in reduced frequencies of caffeine- and fluconazole-resistant colonies. Double mutant *pek1Δ epe1-ΔN150* cells exhibited a similar frequency of caffeine and fluconazole resistant colonies as *epe1-ΔN150* cells. Thus, Pek1 function is bypassed by constitutive Epe1 truncation, confirming that Pek1 acts to promote resistance through Epe1-to-tEpe1 processing (Fig.7g). We conclude that the CIP/MAPK pathway is required to sense and signal the presence of external stresses that can, in part, be overcome by triggering proteasome-mediated Epe1 processing, its loss from chromatin and thus increased heterochromatin-mediated gene repression which confers resistance^13^ (Fig.8).

**Fig. 8:**
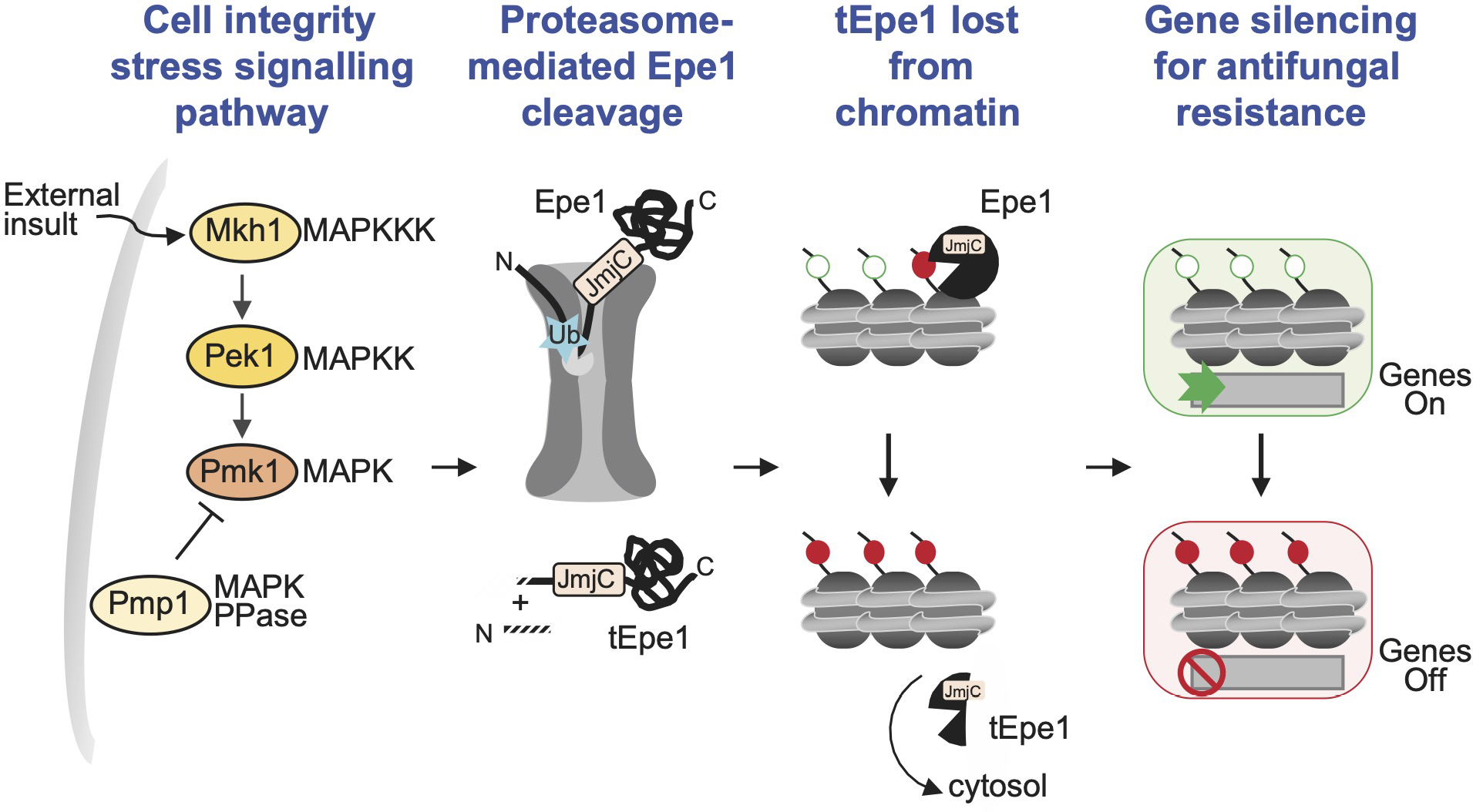
Model for induction of heterochromatin-mediated gene silencing and resulting resistance. External insults (i.e. caffeine, antifungals) detected by the CIP MAPK pathway result in increased Epe1 ubiquitylation (Ub) and proteasome-mediated processing of Epe1 to tEpe1. Loss of the Epe1 N-terminal region results in reduced Epe1 demethylase chromatin association, increased soluble and cytosolic Epe1 and accumulation of Hek9me/heterochromatin over various location, including facultative islands where resulting repression of embedded genes can confer resistance.

## Discussion

We have uncovered an unusual mechanism by which external stresses alter the location of the pivotal heterochromatin regulator Epe1 by inducing its proteasome-mediated processing to a shorter form lacking the N-terminal 150 residues. As previously demonstrated^13^, epimutations arise due to increased heterochromatin formation over genes at various locations, allowing the survival of cells where reduced expression of underlying genes imparts beneficial resistance phenotypes in the face of external challenges. Such genes include *hba1^+^*, encoding a factor involved in nuclear export of the Pap1 stress-induced transcription factor, and *cup1^+^*, encoding a mitochondrial LYR domain protein.

Proteasome-mediated ubiquitin-dependent protein degradation is well known. However, it is less appreciated that some proteins are subject to proteasome-mediated processing in which only part of a target protein is degraded in a process termed regulated <u>u</u>biquitin <u>p</u>roteasome- dependent processing (RUP)^33, 50–58^. Mono-ubiquitylation of target proteins can be sufficient to stimulate such processing events^59, 60^ which frequently terminate at nearby tightly folded structural domains^37, 56^. Thus, elevated Epe1-Myc associated ubiquitylation and enrichment of 23 proteasome subunits with Epe1-GFP upon caffeine treatment (Fig.3a, Supp. Fig.3d) suggest that the translocating activities of the 19S regulatory particle may stall at the JmjC domain. Subsequent cleavage and removal of the N-terminal 150 residues by peptidases within the 20S proteasome core would produce truncated tEpe1 (Fig.8). Proteasome-dependent processing is known to result in ill-defined cleavage sites in substrate proteins due to bidirectional proteasome processivity following initiating cleavage events in the proteasome’s catalytic chamber^37^. Thus, our inability to detect a precise cleavage site is consistent with Epe1 cleavage being mediated by the proteasome. Such proteasome-mediated Epe1 processing may be relevant to the observation that 19S AAA-ATPase mutants, that do not affect ubiquitin-dependent proteolysis, exhibit an altered heterochromatin distribution^61^.

Epe1-to-tEpe1 processing requires an intact cell integrity signalling pathway (CIP/MAPK) to communicate the presence of external insults (Fig.7). Both Pmk1/MAPK and Pek1/MAPKK are needed for stress-induced Epe1 truncation whereas the counteracting MAPK phosphatase Pmp1 acts to restrain tEpe1 generation. We conclude that activation of the CIP/MAPK pathway by external insults results in regulated N-terminal truncation of Epe1 by proteasome-mediated cleavage (Fig.8). CIP/MAPK may directly phosphorylate Epe1 triggering its ubiquitylation. Alternatively, it might act equivalently to MAPK/Slt2^Mpk1^, its *S. cerevisiae* counterpart, which stimulates the assembly of complete 26S proteasomes in response to stress^62^.

Removal of residues 1-150 from Epe1 reduces its chromatin association and nuclear levels, increasing its presence in the cytoplasm, thereby permitting H3K9-methylation, and thus heterochromatin, to increase at known facultative heterochromatin islands and other locations (Fig. 5). Constitutive tEpe1 (Epe1ΔN150) expression also increases the frequency of caffeine and antifungal resistance relative to wild-type cells. The majority of Epe1ΔN150 caffeine-resistant isolates examined exhibited high H3K9me2 levels at *isl14/ncRNA394* (Fig. 6) where reduced expression of the nearby *cup1^+^* encoded mitochondrial protein confers caffeine and anti-fungal resistance^13^. Mitochondrial dysfunction is known to confer antifungal resistance in fission yeast: resistance may result from increased intracellular oxidative stress and associated Pap1 nuclear import, a key stress-regulated transcription factor which elevates efflux pump expression^63–65^. Moreover, higher mitochondrial membrane potential in *S. cerevisiae* correlates with greater anti-fungal resistance; possibly due to efflux pump upregulation^66^. Thus, pathways that respond to mitochondrial-generated stress appear to confer resistance. Interestingly, the JmjC demethylase domain of constitutively truncated Epe1ΔN150 contributes to resistance since mutants expected to disrupt its catalytic activity display reduced resistance (Supp. Fig.6b). Given the increased cytoplasmic presence of Epe1ΔN150 (Fig.4a,d), it is possible that processed tEpe1 acts through unknown cytoplasmic substrates that perhaps alter metabolism and the levels of metabolites that contribute to heterochromatin regulation via chromatin modifiers, thereby influencing resistance^67^.

Constitutive Epe1 truncation causes H3K9me/heterochromatin to redistribute without any selection imposed by external insults. H3K9me2 increases over genes at various locations including heterochromatin islands and especially telomeres, where gene expression is substantially decreased (Fig.5b,d,e; Supp. Fig.5). We therefore propose that in wild-type cells continual exposure to insults triggers Epe1-to-tEpe1 proteasome-mediated processing and selects for the survival of cells where the resulting heterochromatin-imposed gene repression confers resistance through epimutations such as those at *isl14/ncRNA394* that reduce *cup1^+^* expression (Fig.6c)^13^. Insults that trigger Epe1-to-tEpe1 processing include caffeine, salt, hydrogen peroxide and several clinical and agricultural azole-based antifungal compounds (Fig.1b; Fig.6d). Hence fission yeast can transiently adapt to, and resist, external insults by altering the properties and location of Epe1.

The results presented identify a route by which wild-type fungal cells use chromatin-based epigenetic reprogramming to generate epimutations that survive environmental challenges such as those posed by antifungals. Unlike genetic mutants, such resistant epimutants remain wild-type and can thus return to the normal baseline wild-type state upon removal of the triggering insult. However, long-term exposure to that insult will impose selection amongst surviving epimutants for irreversible genetic mutations which fix the resistant phenotype but may also reduce overall fitness. Thus, insult-triggered proteasome-mediated Epe1 processing provides a bet-hedging strategy by which wild-type cells can develop transient resistance with only temporary fitness impacts. Such epigenetic regulation may explain in part the prevalence and variability of resistance in pathogenic plant and animal fungi and may also contribute to the dearth of available effective antifungal treatments.

Azole-based antifungal overuse has resulted in increasing incidence of resistance in both clinical and agricultural settings. H3K9-methylation-dependent heterochromatin is conserved in human pathogenic fungi such as *Cryptococcus*^68^ and *Aspergillus* ^69^, which threaten the health of immunocompromised individuals, and in major crop pathogens such as *Zymoseptoria tritici*^70^ and *Magnaporthe oryzae*^71^ which regularly and significantly reduce global cereal grain yields. Our findings suggest that the development of fungal-specific inhibitors of processes that regulate or mediate heterochromatin formation, and do not affect related host activities, could increase the sensitivity of pathogenic fungi to antifungal compounds. Such compounds would be expected to improve the prognosis of affected patients and reduce crop destruction.

## Acknowledgements

We thank D. Kelly (WCB, Edinburgh) for microscopy and instrumentation support; members of the Allshire Lab for valuable discussions and input; T. Urano for the 5.1.1 (H3K9me2) antibody; A. Fellas for GFP expressing and *clr4*Δ strains; K. Gull for *α*-tubulin antibody; K. Sawin for Mto2 antibody; A.L. Marston for the *Sgo1-GFP S. cerevisiae* strain; J. Svejstrup for provision of the MultiDsk2 expression construct; M.D. Wilson for comments on the manuscript; and colleagues at WCB for support and encouragement during a difficult 2020/21.

## Author Contributions

I.Y. and S.A.W. designed and performed experiments. S.T.-G. contributed to ChIP, ChIP-seq and RNA-seq experiments and CRISPR design. M.L. carried out ChIP-seq and RNA-seq analyses and data visualisation. E.G. performed Halo-tag experiments. R.Y. contributed to antifungal resistance experiments. C.S. ran samples and performed mass spectrometry analysis with J.R. A.L.P. advised on experimental design, performed ChIP and constructed strains. I.Y., S.A.W., A.L.P., and R.C.A. prepared figures and wrote the manuscript.

## Open Access

This research was funded in whole or in part by the Wellcome Trust [Grant numbers 095021, 108504, 200885, 203149, 205008, 218470, and 203149]. For the purpose of Open Access, the author has applied a CC-BY public copyright licence to any author-accepted manuscript version arising from this submission.

## Methods

### Yeast strains and manipulations

*S. pombe* strains used in this study are described in Supp. Table 1. Oligonucleotide sequences are listed in Supp. Table 2. N-terminus truncated Epe1 mutants, 3xMyc-Epe1 and *epe1* deletion strains were constructed by CRISPR/Cas9-mediated genome editing using the SpEDIT system^72^; available on Addgene (#166698, #166699, #166700) with oligonucleotides listed in Supp. Table 2. Epe1-GFP and Epe1-13Myc strains and deletion mutants were generated by using the Bahler tagging and deletion method^73^.

### Protein extraction and western analysis

Cells were treated with 14mM caffeine, 1M KCl, 1mM H2O2 for 16 hours, with fresh H2O2 added every 2 hours to compensate for its decomposition. Protein samples were prepared essentially as previously detailed in Braun, et al, 2011. Briefly, a 10 ml culture of log phase cells was harvested, resuspended in 1 ml of cold water and transferred to an eppendorf tube. Cells were pelleted and resuspended in 775*μ*l water. 150*μ*l 1.85M NaOH and *β*- mercaptoethanol 7.5% were added and the samples incubated on ice for 15 minutes. 150*μ*l 55% TCA was added and incubated for a further 10 minutes on ice. Samples were centrifuged at 15,000 rpm for 10 minutes and pellets were resuspended in HU-buffer (8M urea, 5% SDS, 200mM Tris pH 6·8, 1mM EDTA, with bromo-phenol blue, 1·5 DTT).

Samples were run on a 4-12% NuPAGE Bis-Tris gel (ThermoFisher) and blotted using the MiniBlot Module (Life Technologies) 20V for 1 hour. Western blotting detection was performed using anti-myc 9B11 (Cell Signalling) 1:1000, anti GFP (Roche) 1:1000 and anti-Mouse IgG (whole molecule)–Peroxidase antibody (Sigma) 1:10000. Gels were visualised using the ChemiDoc imaging system (BioRad) and analysed with ImageJ. Where quantified, levels of full-length (FL) Epe1 and truncated tEpe1 were normalised to no treatment and adjusted relative to the *α*-tubulin loading control. Resulting numbers are an average of 3 biological replicates.

### Ubiquitin assay

Harvested cells were washed with PBS and frozen in liquid nitrogen. Frozen cell pellets were ground using a Retsch MM400 mill. Ground cells were resuspended in lysis buffer (10 mM Tris pH7.4, 5 mM CaCl2, 5 mM MgCl2, 50 mM NaCl, 0.1% IGEPAL-CA630, 20 µM MG132, 50 µM PR-619 supplemented with protease inhibitor (Merck Life Sciences) and thawed for 30’ on ice. After centrifugation at high speed, supernatants were subjected to immunoprecipitation with anti-Myc antibody crosslinked to Protein G Dynabeads beads (ThermoFisher) for 2 hrs. After three washes with lysis buffer, beads were resuspended in NuPage LDS sample buffer (ThermoFisher), separated on 4-12% NuPAGE Bis-Tris gel (ThermoFisher) and probed with anti-Myc (9B11; Cell Signalling) and anti-ubiquitin (BML-PW8810; Enzo) antibodies.

### MultiDsk2 Pull Down assay

MultiDsk2 pull-down assays were performed as previously described with slight modifications^74^. Ground cell power was resuspended in D-buffer (150 mM Tris-Acetate pH 7.4, 100 mM potassium acetate, 1 mM EDTA, 0.1% Triton X-100, 10% glycerol, 20 µM MG132 and 50 µM PR-619 Protease Inhibitor (Merck Life Sciences)) and incubated 30’ on ice. After centrifugation, 5 mg of protein from each sample was incubated for two hours with purified Multidsk2 recombinant protein bound to glutathione agarose magnetic beads (Sigma G0924). After incubation beads were washed four times with D-buffer and resuspended in NuPage LDS sample buffer (ThermoFisher) and subjected to western analysis as described above.

### Serial dilution and colony count assays

Equal amounts of cells were grown in YES without agents then serially diluted five-fold and then spotted onto appropriate media. Cells were grown at 30-32°C for 7 days and then imaged. For resistance assays agents were added at the following concentrations; 16 mM caffeine (Merck Life Sciences), 1.8 M KCl, 0.5 mM and 0.75 mM Fluconazole (Merck Life Sciences), 0.0016 mM Tebuconazole (Merck Life Sciences) and 0.006 mM Enilconazole (Merck Life Sciences). 10^4^ cells were plated on 8 plates per strain in triplicate (three independent cultures) biological replicates and resistant colonies that formed after seven to ten days were counted. Significance of the difference between samples was evaluated using Student’s *t*-test. (*) *P* <0.033, (**) *P* <0.002; (***) *P* <0.0002; (****) *P* <0.0001; (n.s). not significant.

### Immunoprecipitation and mass spectrometry analysis

IP and mass spectrometry were performed as previously described^75^. Cell were grown in 4X- YES and 2x10^10^ cells were used per immunoprecipitation experiment. After harvesting, cells were washed twice with water and frozen in liquid nitrogen. Frozen cell pellets were ground using Retsch MM400 mill. Ground cells were resuspended in lysis buffer (10 mM Tris pH7.4, 5mM CaCl2, 5mM MgCl2, 50 mM NaCl, 0.1% IGEPAL-CA630 and supplemented with protease inhibitor (Merck Life Sciences) and 2 mM PMSF (Merck Life Sciences) and thawed for 30 minutes on ice. Chromatin and its associated proteins were solubilized by incubation with 4- 20 units of Micrococcal nuclease (MNase; Merck Life Sciences) for 10 minutes at 37° C. MNase digestion was stopped by adding EGTA to 20 mM and lysates were rotated at 4° C for 1 hour to ensure chromatin solubilization. Supernatant was separated from lysates by centrifugation at 20,000xg for 10 minutes and used for immunoprecipitation using 20μg of anti-GFP antibody (Roche) coupled to 50μl of Protein G Dynabeads (Life Technologies) using dimethyl pimelimidate (DMP; Life Technologies). Bead-bound affinity-selected proteins were washed three times with lysis buffer. Elution was performed with 0.1% RapiGest SF (Waters) in 50mM Tris-HCl pH8 by incubating at 50*°*C for 10 minutes.

RapiGest-eluted samples were denatured by adding final 25mM DTT to the and incubated at 95° C for 5 minutes. Samples were cooled at room temperature and 200μl of 8M Urea in 100mM Tris-HCl pH8 was added and passed through Vivacon 500 column (Sartorius Vivacon 500, 30,000 MWCO Hydrosart) by centrifugation at 14,000xg for 30 minutes. Cysteine residues were alkylated to prevent free sulfhydryls from reforming disulfide bonds by incubation with 100μl of 0.05 M Iodoacetamide (Sigma) in 8 M urea for 20 minutes at 27°C in the dark and centrifuged at 14,000xg to remove the supernatant. Columns were then washed once with 100μl of 8M urea and twice with 100μl of 0.05 M ammonium bicarbonate (Sigma). Proteins were digested by addition of 100μl of 0.05 M ABC containing 0.3 μg of trypsin (90057, Thermo Scientific) at 37°C for 16 hours. Columns were spun at 14,000xg to collect the digested peptides and washed again with 100μl of 0.05 M ABC. Trypsinization was stopped by addition of 10μl of 10% trifluoroacetic acid (TFA) to bring the pH to below 2. C18 reverse-phase resin (Sigma) was used to desalt peptide samples prior to LC-MS/MS. StageTips were packed tightly with 2 layers of C18 resin. The resin was conditioned with 30μl of 100% methanol, washed with 30μl of 80% acetonitrile to remove impurities and finally equilibrated by passing 30μl of 0.1% TFA. The trypsinized peptide solution was passed through the stage tip by centrifugation at 2600 rpm for binding.

Following digestion, samples were diluted with equal volume of 0.1% TFA and spun onto StageTips^76^. Peptides were eluted in 40μl of 80% acetonitrile in 0.1% TFA and concentrated down to 1μl by vacuum centrifugation (Concentrator 5301, Eppendorf, UK). The peptide sample was then prepared for LC-MS/MS analysis by diluting it to 5μl by 0.1% TFA.

LC-MS analyses were performed on an Orbitrap Fusion™ Lumos™ Tribrid™ Mass Spectrometer (Thermo Fisher Scientific, UK) both coupled on-line, to an Ultimate 3000 HPLC (Dionex, Thermo Fisher Scientific, UK). Peptides were separated on a 50 cm (2 µm particle size) EASY-Spray column (Thermo Scientific, UK), which was assembled on an EASY-Spray source (Thermo Scientific, UK) and operated constantly at 50°C. Mobile phase A consisted of 0.1% formic acid in LC-MS grade water and mobile phase B consisted of 80% acetonitrile and 0.1% formic acid. Peptides were loaded onto the column at a flow rate of 0.3 μl min^-1^ and eluted at a flow rate of 0.25 μl min^-1^ according to the following gradient: 2 to 40% mobile phase B in 150 min and then to 95% in 11 min. Mobile phase B was retained at 95% for 5 min and returned back to 2% a minute after until the end of the run (190 min).

Survey scans were recorded at 120,000 resolution (scan range 350-1500 m/z) with an ion target of 4.0e5, and injection time of 50ms. MS2 was performed in the ion trap at a rapid scan mode, with ion target of 2.0E4 and HCD fragmentation^77^ with normalized collision energy of 27. The isolation window in the quadrupole was 1.4 Thomson. Only ions with charge between 2 and 7 were selected for MS2. Dynamic exclusion was set at 60 s.

The MaxQuant software platform^78^ version 1.6.1.0 was used to process the raw files and search was conducted against *Schizosaccharomyces pombe* complete/reference proteome set of PomBase database (released in July 2017), using the Andromeda search engine ^79^ For the first search, peptide tolerance was set to 20 ppm while for the main search peptide tolerance was set to 4.5 pm. Isotope mass tolerance was 2 ppm and maximum charge to 7. Digestion mode was set to specific with trypsin allowing maximum of two missed cleavages. Carbamidomethylation of cysteine was set as fixed modification. Oxidation of methionine, phosphorylation of serine, threonine and tyrosine and ubiquitination of lysine were set as variable modifications. Additionally, methylation of lysine and arginine, di- and tri-methylation of lysine were also set as variable modifications. Label-free quantitation analysis was performed by employing the MaxLFQ algorithm as described^80^. Absolute protein quantification was performed as described previously^81^. Peptide and protein identifications were filtered to 1% FDR.

### Cytology

Cells were fixed with 3.7% formaldehyde (Sigma) for 10 minutes at room temperature. Immunolocalization staining was performed as described previously^82^. The following antibodies were used; anti-CENP-A^Cnp1^ (sheep in-house, 1:3000), anti-GFP (ThermoFisher Scientific, 1:200), Alexa 594 and 488 labelled secondary antibodies at 1:1000 dilution (Life Technologies). Images were acquired with Zeiss Axioplan, using a 100X/1.40 NA Plan-Apochromat Oil DIC M27 objective lens and processed using Metamorph acquisition and processing software (Zeiss MicroImaging) or FIJI.

For imaging cells with Halo tagged proteins, 250 to 500nM of HaloTag ligand JF646 dye was added to 400*μ*l of log phase cells and incubated at 32°C for 3 hours. Cells were washed with PMG and plated on a PMG agarose pad, dried before addition of a coverslip. Images were taken with an inverted microscope (Nikon TiE) equipped with an EMCCD Camera (iXion Ultra 897, Andor), a SpectraX Line engine (Lumencor) and a 100X Nikon objective (NA 1.45, oil immersion). For HaloTag ligand JF646, a red LED and a CY5 filter cube were used. Images were taken with an EM Gain of 300 and exposure times between 20 and 200ms per frame.

For all colour channels, a series of 11 z slices (step size 0.2 μm) were taken to cover the entire depth of the cells. Live cells were kept at 32°C during imaging. All image analysis was performed in FIJI as follows: a region of interest was manually selected (nuclear/cytoplasmic spot). Background intensity for each cell was subtracted from the mean intensity measurement. Significance of the difference between samples was evaluated using Student’s *t*-test. (*) *P* <0.0.33, (**) *P* <0.002; (***) *P* <0.0002; (****) *P* <0.0001; (n.s). not significant.

### qRT-PCR

Total RNA was extracted using 5x10^7^ cells and Monarch Total RNA Miniprep Kit (NEB) as per manufacturer’s instructions. Samples were treated with Turbo DNase (Ambion) to remove contaminating DNA. Reverse transcription was performed using LunaScript RT SuperMix Kit (NEB). Oligonucleotides used for qRT-PCR are listed in Supp. Table 1. qRT-PCR histograms represent 3 technical replicates; error bars correspond to standard error of the mean.

### Fractionation

Cells were grown in 4xYES and harvested by centrifugation at 3500×g, washed twice with water and flash frozen in liquid nitrogen. Frozen cell pellets were ground using Retsch MM400 mill. Ground cells were resuspended in lysis buffer (10mM Tris pH7.4, 5mM CaCl2, 5mM MgCl2, 50mM NaCl, 0.1% IGEPAL-CA630 and supplemented with protease inhibitor (P8215, Sigma) and 2 mM PMSF After thawing on ice for 30 minutes, sample were vortexed 15 times for 30”, with one minute on ice between each wash. After centrifugation at 5,000xg, supernatant was collected as soluble fraction, pellet was washed twice with same lysis buffer and then collected as insoluble fraction.

### ChIP

ChIP experiments were performed as previously described^83^ using anti-H3K9me2 (5.1.1, a kind gift from Takeshi Urano) or anti-GFP (Life Technologies). Immunoprecipitated DNA was recovered with Chelex-100 resin. qChIPs were analysed by real-time PCR using Lightcycler 480 SYBR Green (Roche) Oligonucleotides are listed in Supp. Table 1. ChIP enrichments were calculated as % DNA immunoprecipitated at the locus of interest relative to the corresponding input samples. For H3K9me2 ChIP, formaldehyde fixed *Schizosaccharomyces octosporus* cells were added at the cell lysis stage for spike-in control. For Epe1-GFP ChIP, formaldehyde-fixed Sgo1-GFP S*accharomyces cerevisiae* cells^84^ were added at the cell lysis stage for spike-in control. Histograms represent data averaged from three biological replicates; error bars represent standard deviations. Significance of the difference between samples was evaluated using Student’s *t*-test. (*) *P* <0.0.33, (**) *P* <0.002; (***) *P* <0.0002; (****) *P* <0.0001; (n.s). not significant.

### ChIP-seq, RNA-seq and Bioinformatic analyses

Samples were prepared as previously described^13^. ChIP-seq and RNA-seq samples were compared using DESeq2 statistics of read counts per feature, accessed by HTSeq-count and genomic 1kb bins for ChIP-seq or by Salmon-quant and annotated genes for RNA-seq analyses. Features with adjusted p-values <0.1 were selected as significantly changed. Features with log2 (Fold Change) values >1 (RNA-seq) or >2 (ChIP-seq) are presented in figures.

## Supplementary Information

List

Supplemental Fig 1 and legend

Supplemental Fig 2 and legend

Supplemental Fig 3 and legend

Supplemental Fig 4 and legend

Supplemental Fig 5 and legend

Supplemental Fig 6 and legend

Supplemental Fig 7 and legend

Supplemental Table 1 – available as Excel file (Dataset 1) on request

Supplemental Table 2 – available as Excel file (Dataset 2) on request

Supplemental Table 3 – available as Excel file (Dataset 3) on request

Supplemental Table 4 – available as Excel file (Dataset 4) on request

Supplemental Table 5 – available as Excel file (Dataset 5) on request

Supplemental Table 6 – available as Excel file (Dataset 6) on request

Supplemental Table 7 – available as Excel file (Dataset 7) on request

## Supplementary Figures and Legends

**Supplemental Fig. 1.**
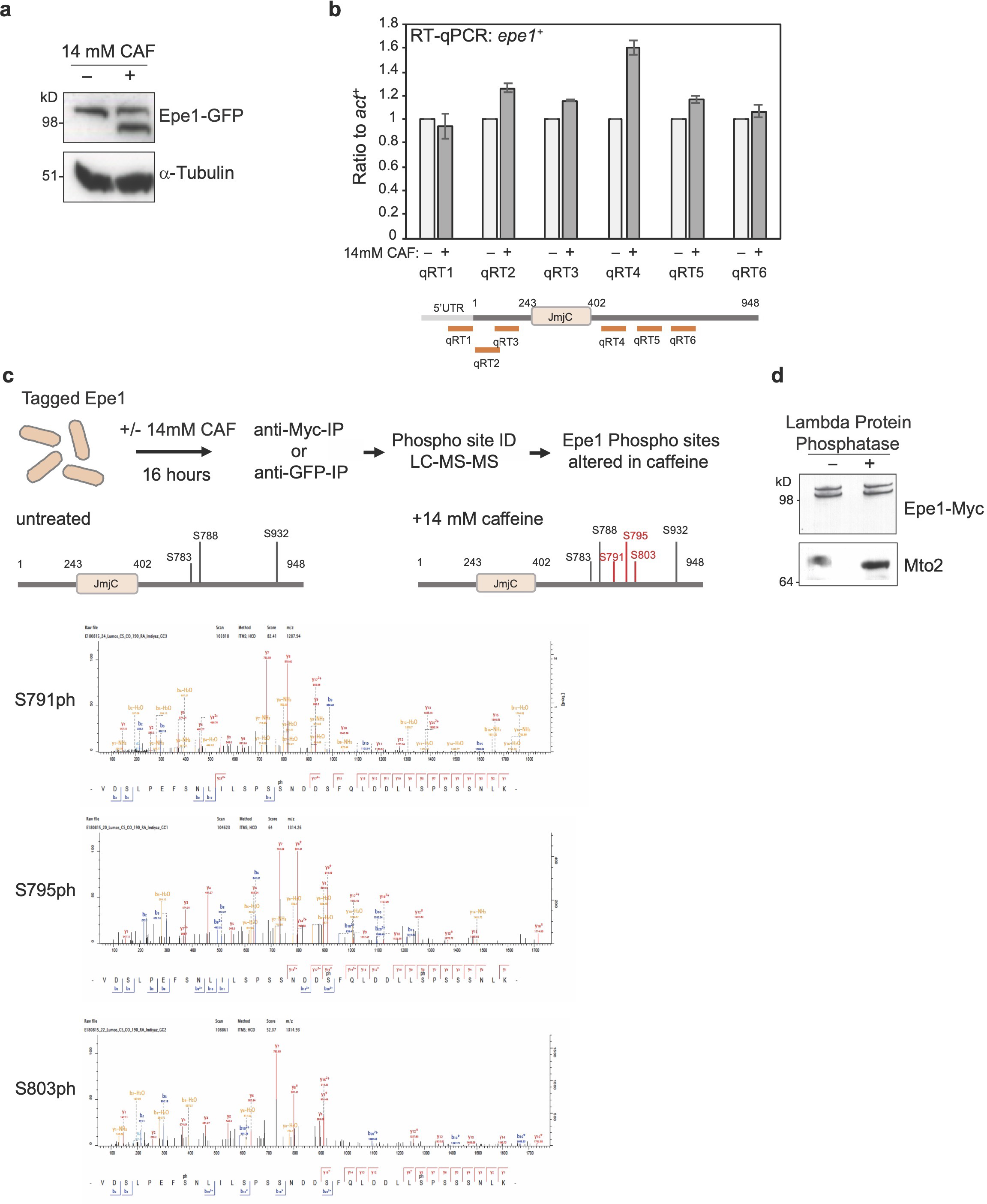
Caffeine-induced alteration in Epe1-GFP mobility does not result from transcriptional changes in 5’ region of *epe1^+^* gene or Epe1-Myc phosphorylation. a. Epe1-GFP or *α*-tubulin western from cells untreated (-)/treated (+) with 14mM caffeine for 16h. b. qRT-PCR measurement of steady-state Epe1 transcript levels in cells untreated (-)/treated (+) with 14mM caffeine for 16h. Locations of primers used indicated. c. Scheme to detect caffeine-induced Epe1-GFP phosphorylation by mass spectrometry (top). Representation of phosphorylated residues detected on Epe1-GFP from cells untreated or treated with 14mM caffeine for 16 h (middle). Spectra showing detection of phosphorylated serine S791, S795 and S803 residues after caffeine treatment (bottom). Full data, Supplementary Table 4. d. Epe1-Myc and Mto2 westerns from cells treated with 14mM caffeine for 16h before (-)/after (+) lambda protein phosphatase addition. Mto2 mobility is known to be Lambda Phosphatase sensitive^1^.

**Supplemental Fig. 2.**
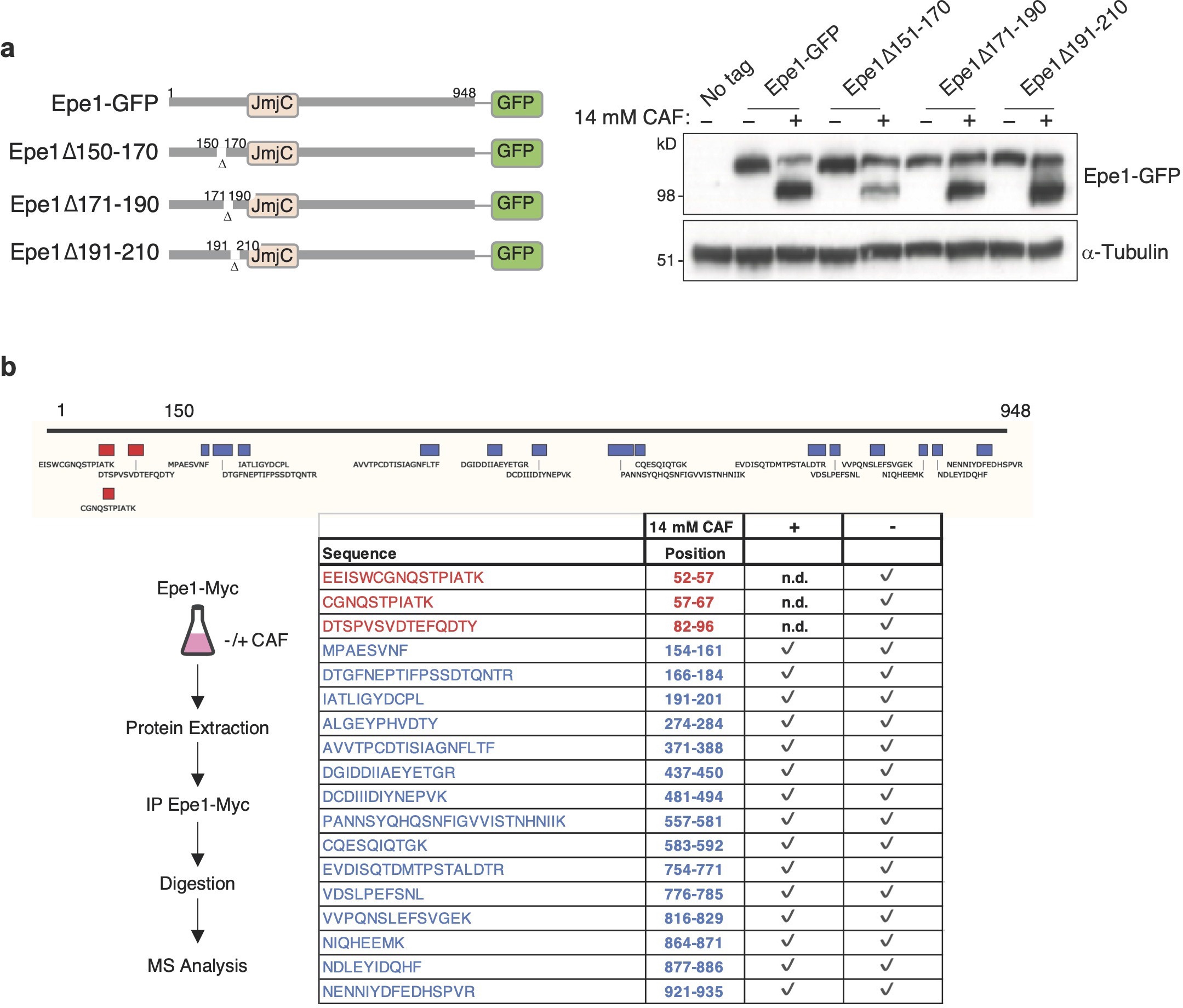
Deletions beyond residue 150 in Epe1-GFP do not hinder caffeine-induced and peptides within the first 100 residues are not detected following caffeine treatment. a. Schematic of indicated 20 residue deletion mutants in the N-terminal coding region of the endogenous *epe1* gene expressed as GFP fusions (Epe1Δ150-170-GFP, Epe1Δ171-190- GFP, Epe1Δ191-210-GFP; left). Western detecting indicated mutant Epe1-GFP fusion proteins or *α*-tubulin from cells untreated (-) or treated (+) with 14 mM caffeine; right). b. Epe1-Myc peptides detected following immunoprecipitation from cells untreated (-) or treated (+) with 14mM caffeine and analysis by mass spectrometry. Top: schematic showing position of peptides detected relative to Epe1 (residues 1-948). Bottom: Epe1 peptides detected in Epe1-Myc immunoprecipitates from treated (+) or untreated (-) with 14mM caffeine. Of the eighteen peptides detected from Epe1-Myc in untreated (-) samples three were not detected (n.d.; red) in the caffeine-treated sample. The three peptides not detected in the presence of caffeine are derived from within the first 100 residues on the N-terminus. Analysis was performed on three independent immunoprecipitates. Full data, Supplementary Table 5.

**Supplemental Fig. 3.**
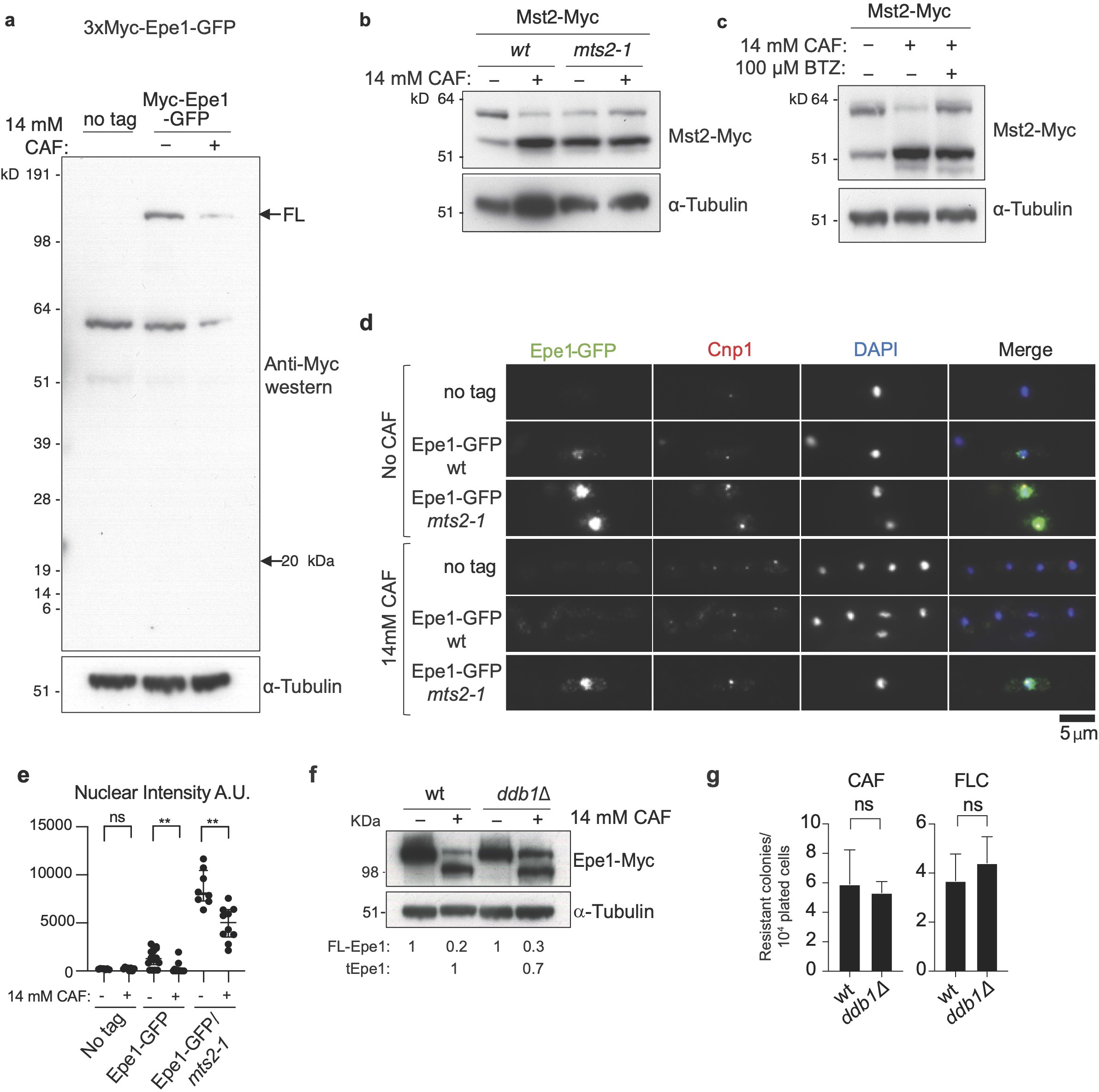
A caffeine-induced N-terminal processing product is not detectable but Epe1-associated ubiquitination increases and Epe1 processing is sensitive to loss of specific E3 ligases which influence resistance. a. Anti-Myc western of cells untreated (-)/treated (+) with 14 mM caffeine showing that a predicted N-terminal 20 kDa Myc-Epe11-150 processing product is not detectable. b. Western showing detection of both isoforms of the histone acetyltransferase Mst2-Myc or *α*-tubulin from wild-type or *mst2-1* proteasome defective cells untreated (-)/treated (+) with 14mM caffeine for 16h. This control demonstrates that other known stress-induced changes still occur in *mst2-1* cells. c. Western showing detection of both isoforms of the histone acetyltransferase Mst2-Myc or *α*-tubulin from wild-type cells incubated without (-)/with (+) with the 26S proteasome inhibitor Bortezomib (BTZ) prior to no caffeine (-)/14mM caffeine (+) treatment for 16h. This control demonstrates that other known stress-induced changes still occur in the presence of BTZ. d. Immunolocalization of Epe1-GFP in wild-type or *mts2-1* cells untreated (-)/ treated(+) with 14 mM/16h caffeine. No tag, negative control. e. Quantification of anti-GFP/Epe1-GFP nuclear signals of cells in d. f. Western detecting Epe1-GFP or *α*-tubulin in wild-type or *ddb1Δ,* cells untreated (-)/treated (+) with 14mM caffeine for 16h. Numbers below tracks: levels of full-length FL-Epe1 and truncated tEpe1 normalised to no treatment and adjusted relative to *α*-tubulin loading control, measurement average of 2 biological replicates. g. Number of resistant colonies formed/1x10^4^ viable cells by wild-type or *ddb1Δ* cells plated on caffeine or fluconazole plates. Statistical analysis of three biological replicates performed as in Materials and Methods.

**Supplemental Fig. 4.**
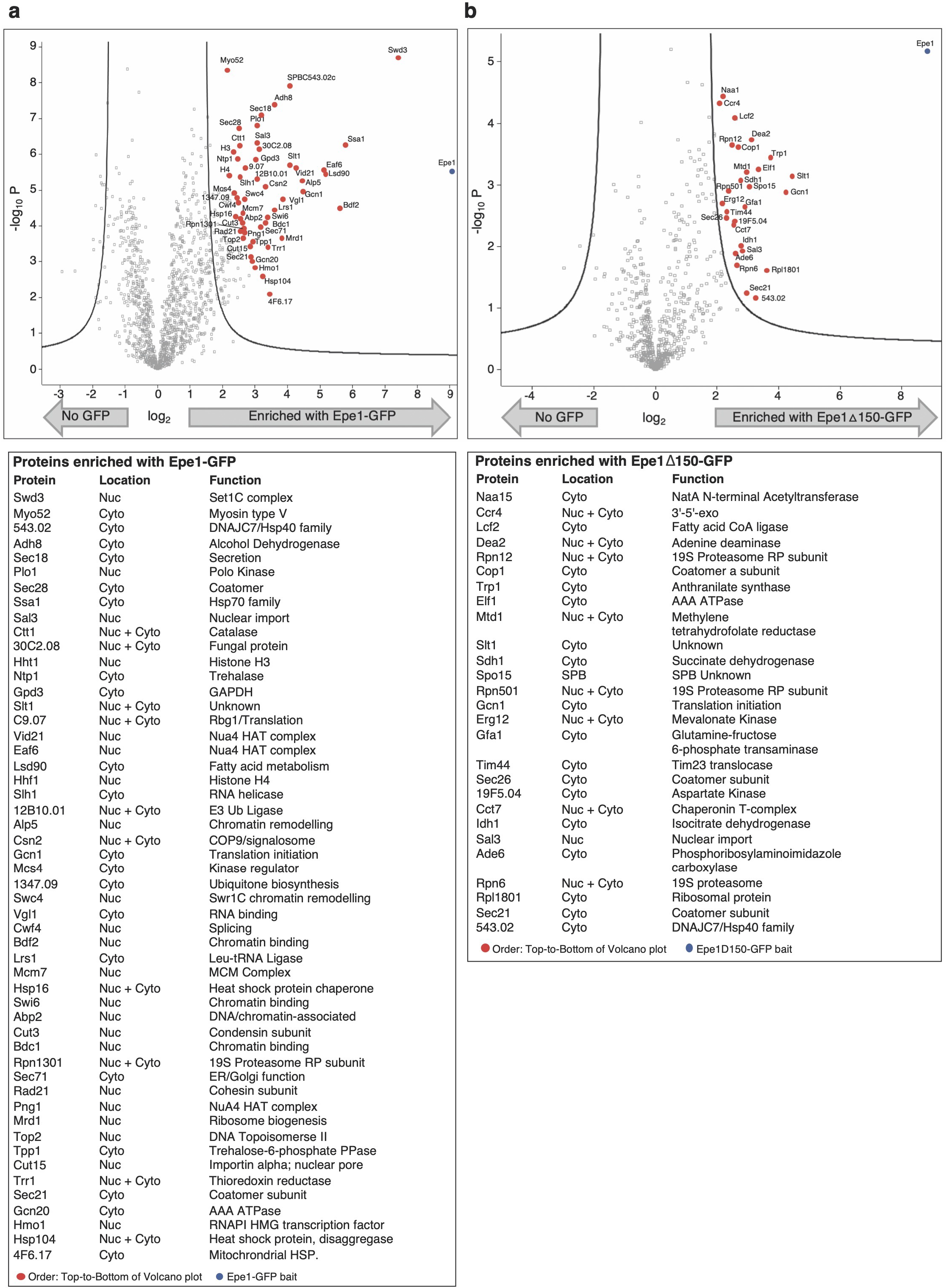
Chromatin-associated nuclear proteins are enriched with full length Epe1 whereas more cytoplasmic proteins are enriched with constitutively truncated Epe1ΔN150-GFP. a. Volcano plot of proteins enriched with full-length Epe1-GFP extracted from cells detected by proteomic analysis (top), specific proteins labelled (red dots). Table with names and functions of proteins enriched (bottom). Full data, Supplementary Table 6. b. Volcano plot of proteins enriched with constitutively processed Epe1ΔN150-GFP extracted from cells and detected by proteomic analysis (top), specific proteins labelled (red dots). Table with names and functions of proteins enriched (bottom). Full data, Supplementary Table 7.

**Supplemental Fig. 5.**
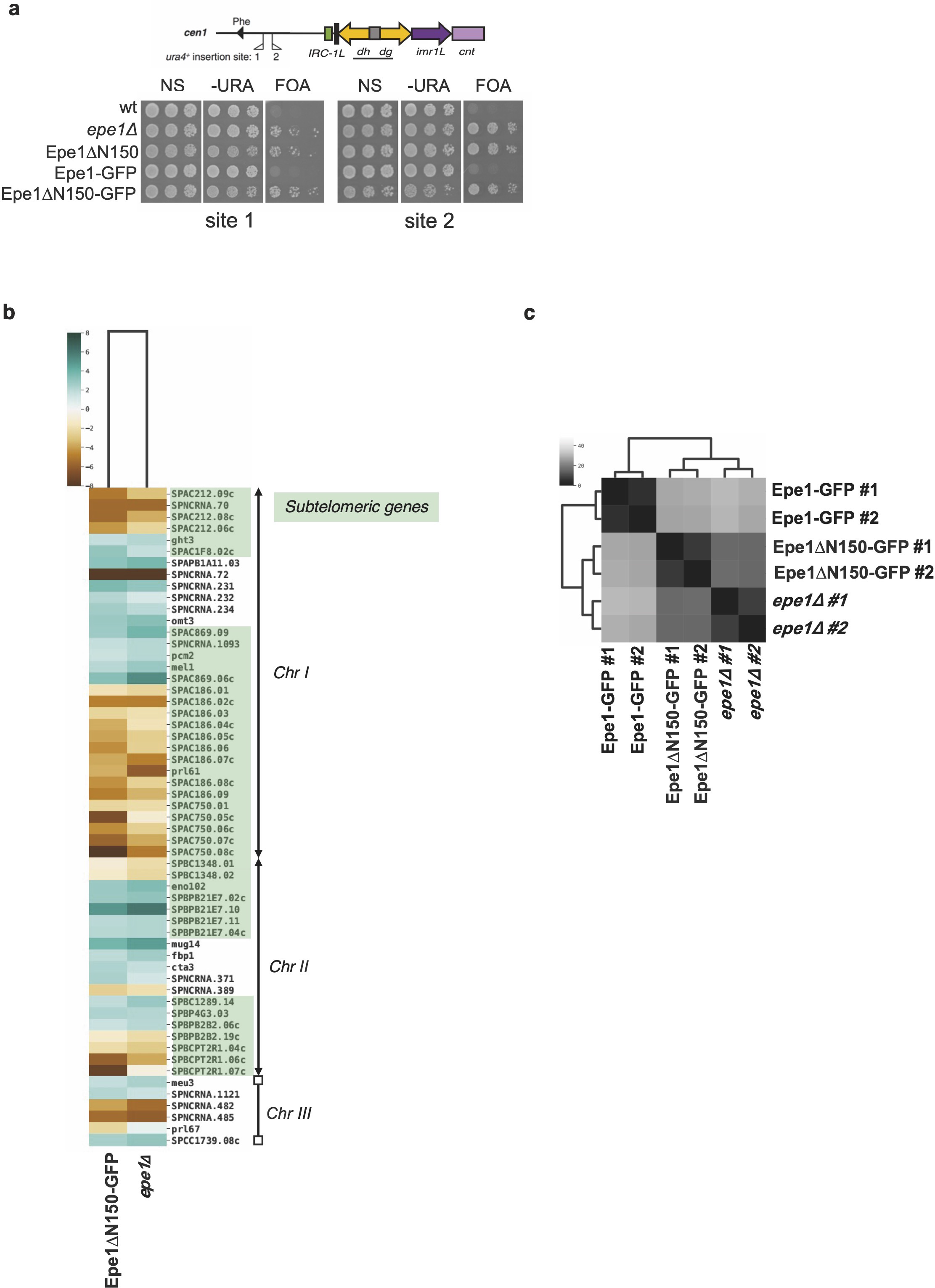
Altered silencing and gene expression in *epe1Δ* and Epe1- ΔN150-GFP cells. a. Wild-type, *epe1*Δ, Epe1-ΔN150, Epe1-GFP/wt and Epe1-ΔN150-GFP cells harbouring *ura4^+^* marker gene insertions at either site 1 or site 2 as indicated (top) were serially diluted and plated on non-selective (N/S), selective (-URA) or counter-selective (FOA) plates. b. Heat-map generated from processed data of RNA-seq two biological replicates from Epe1ΔN150-GFP or *epe1Δ* cells compared with that of wild-type Epe1-GFP cells. Genes with higher (turquoise) or lower (brown) expression relative to wild-type Epe1-GFP cells are shown as a Log2 scale. The position of specific affected genes along *S. pombe* chromosomes Chr I, II, III are shown with arrowheads indicating telomeres on Chr I and ChrII. The annotated *S. pombe* genome shows only rDNA arrays (rectangles) at both ends of Chr III. c. Heat-map showing that RNA-seq data (used in b.) resulting from two Epe1ΔN150-GFP and *epe1Δ* biological replicates are more similar to each other than they are to wild-type Epe1-GFP cells.

**Supplemental Fig. 6.**
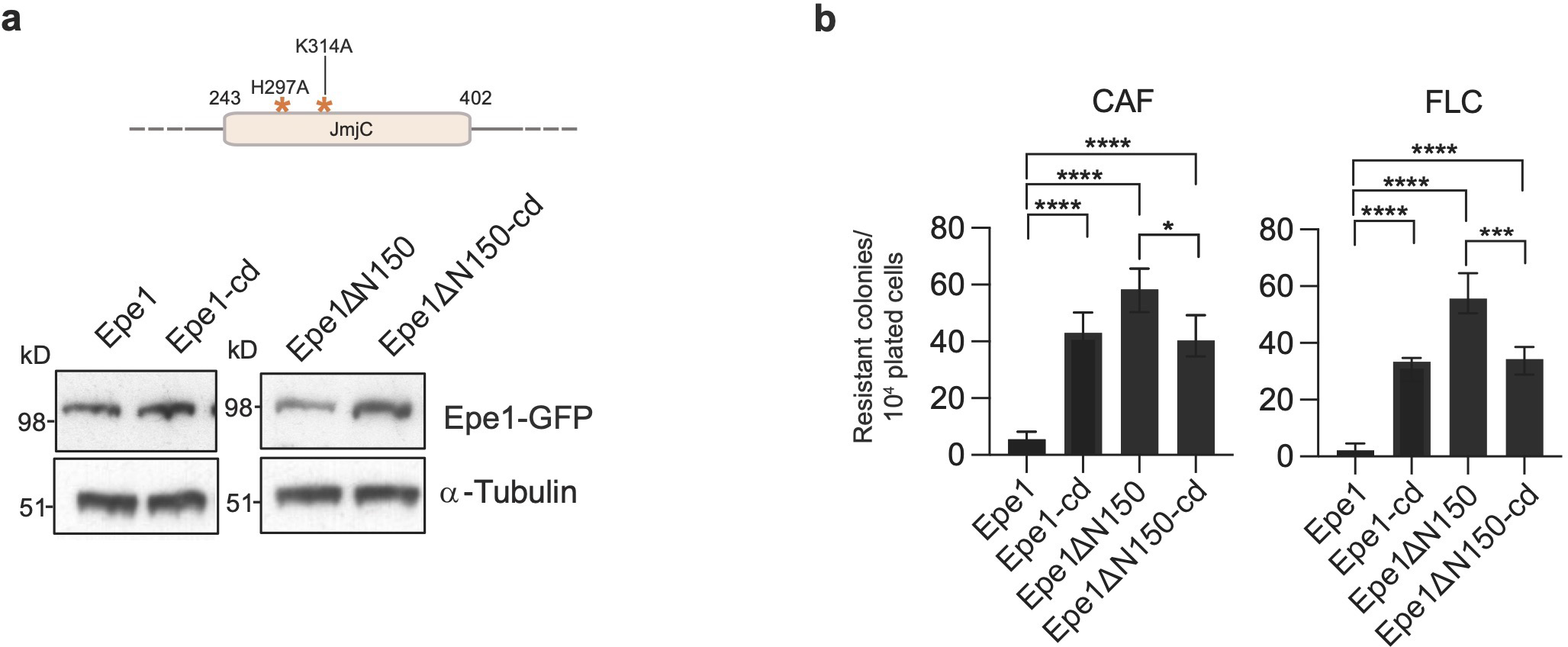
Cells expressing predicted catalytically inactive Epe1-GFP and Epe1-ΔN150-GFP exhibit decreased resistance. a. Schematic showing position of H297A and K314A mutations predicted to reduce association of the essential iron/Fe(2) and 2-Oxoglutarate *(α*-ketoglutarate) cofactors, respectively, with the JmjC demethylase domain of Epe1 (Top). Westerns detecting Epe1- GFP, Epe1-cd-GFP, Epe1ΔN150-GFP and Epe1ΔN150-cd-GFP (cd; catalytically dead) or *α*- tubulin (bottom). b. Number of resistant colonies formed/1x10^4^ viable cells by Epe1-GFP, Epe1-cd-GFP, Epe1ΔN150-GFP and Epe1ΔN150-cd-GFP cells plated on caffeine or fluconazole plates.

**Supplemental Fig. 7.**
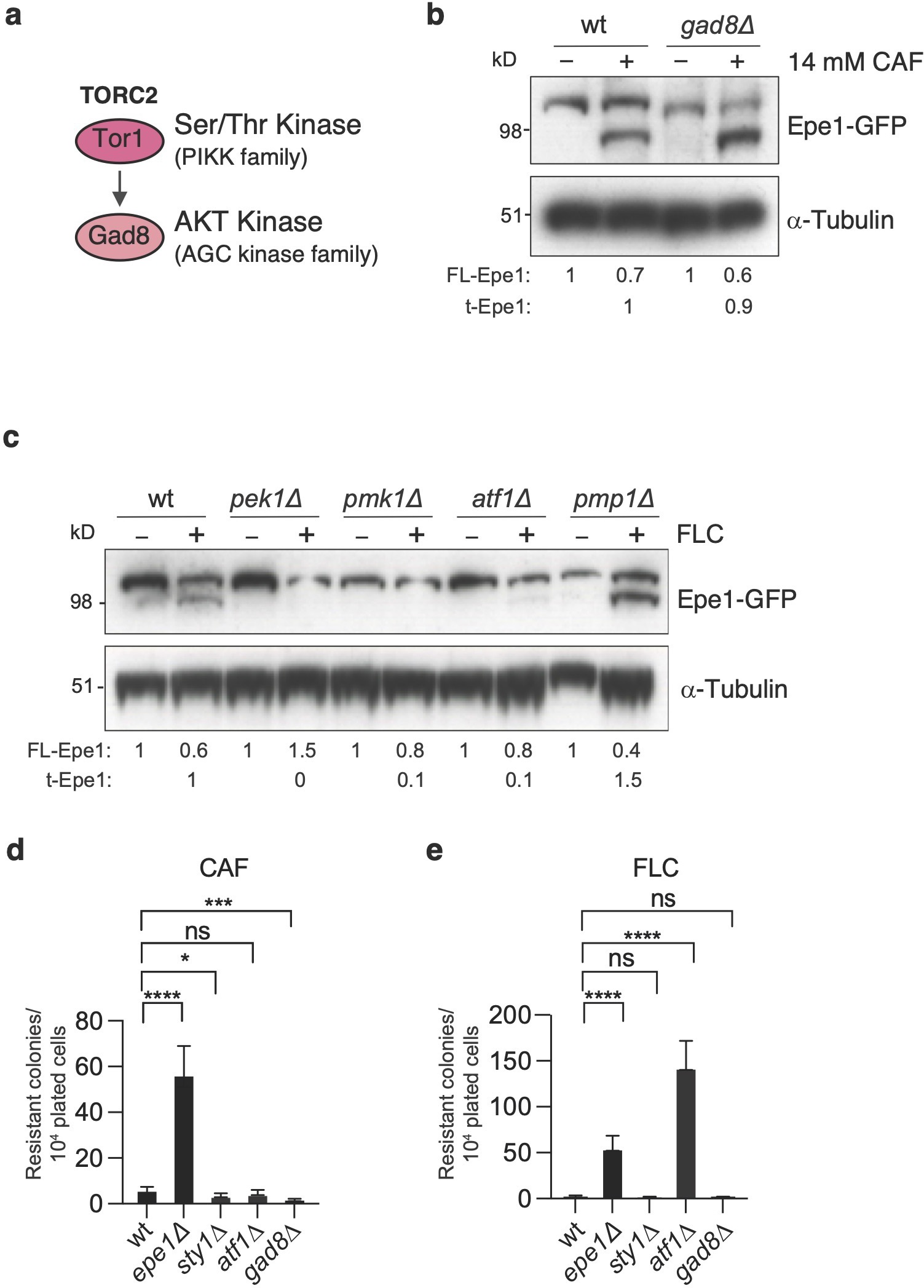
The cell integrity/Pmk1 but not the TORC2/Gad8 stress signalling pathway regulates Epe1 processing and resistance. a. Diagram of part the *S. pombe* TORC2/Tor1-dependent signalling pathway. b. Western detecting Epe1-GFP or *α*-tubulin from wild-type, or *gad8Δ* cells untreated (-)/ treated (+) with 14mM/16h caffeine. c. Western detecting Epe1-Myc or *α*-tubulin from wild-type, *pek1Δ*, *pmk1Δ, atf1Δ* or *pmp1Δ* cells untreated (-)/treated (+) with 0.5mM/16h fluconazole (FLC). d. Number of resistant colonies formed/1x10^4^ viable cells plated by wild-type, *epe1Δ, sty1Δ*, *atf1Δ* and *gad8Δ* on plates containing 16mM caffeine (CAF). e. Number of resistant colonies formed/1x10^4^ viable cells plated by wild-type, *epe1Δ, sty1Δ*, *atf1Δ* and *gad8Δ* on plates containing 0.5mM fluconazole (FLC). Statistical analysis of three biological replicates performed as in Materials and Methods.

